# Engineered illumination devices for optogenetic control of cellular signaling dynamics

**DOI:** 10.1101/675892

**Authors:** Nicole A. Repina, Thomas McClave, Xiaoping Bao, Ravi S. Kane, David V. Schaffer

## Abstract

Spatially and temporally varying patterns of morphogen signals during development drive cell fate specification at the proper location and time. However, current *in vitro* methods typically do not allow for precise, dynamic, spatiotemporal control of morphogen signaling and are thus insufficient to readily study how morphogen dynamics impact cell behavior. Here we show that optogenetic Wnt/β-catenin pathway activation can be controlled at user-defined intensities, temporal sequences, and spatial patterns using novel engineered illumination devices for optogenetic photostimulation and light activation at variable amplitudes (LAVA). The optical design of LAVA devices was optimized for uniform illumination of multi-well cell culture plates to enable high-throughput, spatiotemporal optogenetic activation of signaling pathways and protein-protein interactions. Using the LAVA devices, variation in light intensity induced a dose-dependent response in optoWnt activation and downstream Brachyury expression in human embryonic stem cells (hESCs). Furthermore, time-varying and spatially localized patterns of light revealed tissue patterning that models embryonic presentation of Wnt signals *in vitro*. The engineered LAVA devices thus provide a low-cost, user-friendly method for high-throughput and spatiotemporal optogenetic control of cell signaling for applications in developmental and cell biology.

## INTRODUCTION

Cell fate decisions are governed by dynamic, spatially and temporally varying signals from the cellular environment. In particular, during development, morphogen gradients orchestrate the dynamic, coordinated movement and differentiation of embryonic cell populations (Arnold & Robertson 2009). Genetic perturbation and biomolecular treatment with pathway agonists or inhibitors have provided insight into the critical regulators of embryogenesis, yet spatially-varying interactions between cell subpopulations and time-varying signal dynamics and duration thresholds remain largely unstudied due to a lack of tools to perturb signaling dynamics in biological systems (Arnold & Robertson 2009; Oates et al. 2009).

Optogenetic methods aim to address this need for techniques that control such cell signal dynamics. With optogenetics, light-responsive proteins from plants, algae, and bacteria have been adapted to control signaling and protein-protein interactions in mammalian cells (Boyden et al. 2005; Nagel et al. 2003; Hegemann & Nagel 2013). In particular, a variety of photosensory domains have been discovered, optimized, and repurposed to place intracellular signaling pathways under light control, capabilities that offer the unique advantage of using light patterns to stimulate signaling in a specific location and at a specific time (Repina et al. 2017). Within mammalian cells, optogenetic tools that control protein-protein interactions have for example been developed for diverse signaling pathways such as Wnt/β-catenin (Bugaj et al. 2013; Repina et al. 2019), Ras/Raf/Mek/Erk (Toettcher et al. 2011; Johnson & Toettcher 2019; Toettcher et al. 2013; Johnson et al. 2017), fibroblast growth factor (FGF) (Kainrath et al. 2017), and Rho-family GTPase signaling (Levskaya et al. 2009; Yazawa et al. 2009; Strickland et al. 2012; Guntas et al. 2015; Bugaj et al. 2015; Wang et al. 2010). Optogenetic methods have also been recently applied to studies in developmental biology in key model organisms (Johnson & Toettcher 2018). For example, in *Drosophila* and zebrafish embryos, dynamic control of signaling elucidated the spatiotemporal limits of ERK (Johnson et al. 2017; Johnson & Toettcher 2019), Nodal (Sako et al. 2016), and Bicoid (Huang et al. 2017) patterning and how local modulation of cell migration and contractility drives tissue morphogenesis (Izquierdo et al. 2018; Guglielmi et al. 2015; Čapek et al. 2019). Furthermore, in embryonic stem cell models for mammalian development, we have recently achieved optogenetic control of canonical Wnt signaling and elucidated mechanisms of tissue self-organization during mesendoderm differentiation (Bugaj et al. 2013; Repina et al. 2019).

To activate the variety of photosensory domains developed for cellular signaling pathway control, there must be complementary development of optical tools for cell culture illumination. However, current optical methods lack practical applicability to routine cell culture stimulation and/or lack critical characterization and functionality. In particular, microscope-based systems are widely used for optogenetic photostimulation that implement one- or two-photon excitation to scan a single diffraction-limited spot (Packer et al. 2013; Prakash et al. 2012; Packer et al. 2015; Carrillo-Reid et al. 2016; Nikolenko et al. 2007) or project multi-spot light patterns (Papagiakoumou et al. 2010; Pégard et al. 2017; Papagiakoumou et al. 2008; Hernandez et al. 2016) onto the biological sample. While such systems are essential for high-resolution *in vivo* experiments and precise manipulation of neural circuits, their application to intracellular signal pathway activation in cell cultures is limited by the system complexity, low throughput, high cost, and need for continuous environmental control (Yizhar et al. 2011). For manipulating signaling pathways where speed and single-cell spatial resolution are often less critical, the paramount photostimulation criteria are high-throughput control of multiple parallel biological conditions, defined illumination parameters, and compatibility with established cell culture formats and assays. Several methods have been developed for such photostimulation of signal pathways in cell cultures, though they can lack critical characterization and functionality. For example, a simple panel of light emitting diodes (LEDs) enables optogenetic activation of a tissue culture plate, but lacks multi-well, high-throughput control (Müller et al. 2014; Shao et al. 2018) and spatiotemporal patterning capability (Tucker et al. 2014). Multi-well control can be achieved by incorporating a microcontroller and LED drivers that set user-defined intensities for separate wells (Olson et al. 2014; Gerhardt et al. 2016). Though such multi-well devices have advanced the throughput of optogenetic studies, current implementations have not characterized key performance parameters such as uniformity of illumination, spatial or temporal light patterning resolution, and quantification of device overheating and cell toxicity (Gerhardt et al. 2016; E. A. Davidson et al. 2013; Lee et al. 2013; Hannanta-Anan & Chow 2016; Bugaj et al. 2018; Hennemann et al. 2018; Richter et al. 2015; Olson et al. 2014). Several such designs also rely on modification of expensive instrumentation (Richter et al. 2015) or focus on bacterial culture tubes (Olson et al. 2014).

We have designed a programmable illumination system for photostimulation of multi-well plates that can be readily incorporated into the workflow of routine cell culture and allow controlled and quantitative spatiotemporal light patterning. Specifically, we engineer cell culture illumination devices for light activation <u>a</u>t <u>v</u>ariable <u>a</u>mplitudes, or LAVA boards. We optimize the LAVA board optical configuration for illumination uniformity and achieve programmable photostimulation of independent wells of 24-well or 96-well culture plates kept in standard 37°C tissue culture incubators. Each well can be wirelessly programmed through a graphical user interface (GUI) at user-defined intensities (0 – 20 µWmm^-2^, 0.005 µWmm^-2^ resolution), temporal sequences (10 ms resolution), and spatial patterns (100 µm resolution). We demonstrate LAVA board performance by modulating the intensity, timing, and spatial location of canonical Wnt/β-catenin signaling in human embryonic stem cell (hESC) cultures using the optoWnt optogenetic system (Bugaj et al. 2013; Repina et al. 2019). We show that Wnt pathway activation and hESC differentiation is dose-responsive to light intensity and duration of illumination, and that spatial patterning can be used to simulate the embryonic, spatially polarized presentation of the Wnt ligand. Lastly, we provide a detailed protocol for LAVA board assembly, which takes ∼8 hrs and less than $500 to fabricate and build.

## DESIGN

### Design requirements for spatiotemporal photostimulation of cell cultures

A number of design considerations should be considered to ensure controlled, long-term illumination of mammalian cell cultures. The primary requirement is illumination of cells with defined intensities of light and with illumination patterns that can vary in space and time. Also, the intensity range should be sufficient to activate the photosensory protein of interest, which can vary widely from continuous illumination at 1 µWmm^-2^ (Repina et al. 2019) to short pulses at 1,000 – 10,000 µWmm^-2^ (Yizhar et al. 2011). Moreover, the required temporal resolution also depends on the photosensory domain and experimental application: 1-100 ms pulses are typically used for control of neural circuits (Yizhar et al. 2011), whereas signal pathway dynamics are controlled on the second to multi-day timescales (Repina et al. 2019; Johnson & Toettcher 2018). Furthermore, for spatial resolution requirements, optimal resolution for subcellular stimulation would be 0.2 – 2µm, but such resolutions are infeasible without complex optical systems. Since a typical size of single cells is 10 – 20 µm, multicellular resolution can be sufficient for many applications at 50 – 200 µm.

Beyond the requirement for light delivery, the illumination device must also be compatible with routine stem cell culture experiments. As a result, the electronics must be compatible with the warm (37°C) and humid environment used for mammalian tissue culture. Additionally, the device should also not induce cell damage from overheating or phototoxicity, which is a significant concern during optogenetic stimulation since such processes alter cell physiology and/or cause non-specific activation of signaling pathways (Tyssowski & Gray 2019; Acker et al. 2016; Allen et al. 2015; Yizhar et al. 2011; Owen et al. 2019). Illumination uniformity across a region of interest is critical as well. Since the strength of induced signaling is dependent on light dosage, nonuniformity can give rise to signal variation across the region of optogenetic stimulation. Lastly, cost, functionality, and ease of use can be significant barriers for optogenetic studies, so simple user-programmable control of many simultaneous illumination conditions is a significant advantage for complex and high-throughput biological experiments.

### Design overview of engineered LAVA illumination devices

To enable precise control over the intensity, timing, and location of optogenetic stimulation, we engineered illumination devices, LAVA boards, that incorporate into the workflow of stem cell culture (Figure 1A). LAVA boards project user-defined, programmable light patterns onto 24-well or 96-well tissue culture (TC) plates maintained inside a standard TC incubator (Figure 1B-1C, Figure S1A-S1C). In brief, light emitted by LEDs located underneath the multiwell culture plate passes through diffusive optical elements that ensure uniform illumination of each well. For stimulation of cells expressing optoWnt, a system we previously engineered for optogenetic control of Wnt signaling(Bugaj et al. 2013; Repina et al. 2019), we chose blue LEDs emitting at a central wavelength of 470 nm to match the *A. thaliana* Cryptochrome 2 (Cry2) photosensory domain absorption spectrum (Figure S1D), though we note that LEDs of diverse colors can readily be substituted for use with other optogenetic systems. The LAVA board electronics are designed for programmable control of illumination intensity and temporal sequences, with independent control of each well. In addition, spatial precision is conveyed through an intensity mask attached to the culture plate. The hardware design also includes a cooling system and vibration isolation to ensure cell viability. Lastly, for ease of use, we developed a graphical user interface (GUI) to wirelessly program the illumination intensity and temporal sequences for each well.

**Figure 1.**
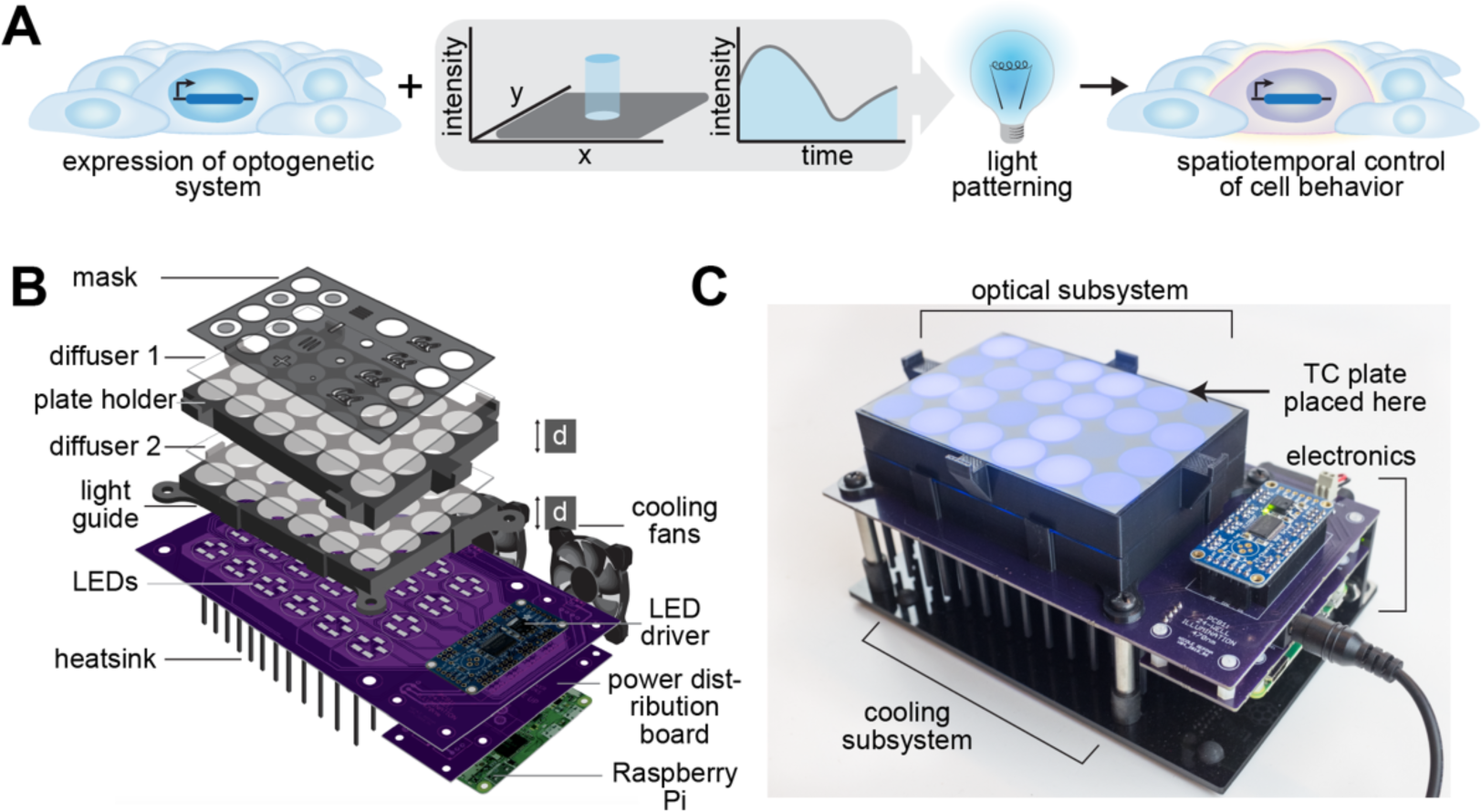
Overview of illumination device, LAVA, for optogenetic stimulation of hESC cultures. **a)** Schematic of typical optogenetic experiment, where spatiotemporal precision is conferred through light patterning **b)** Diagram of illumination device design. LEDs illuminate a TC plate placed on top of device, with light passing through a series of two light guides, two optical diffusers, and a die-cut mask. LEDs are programmed through a Raspberry Pi and LED driver and are cooled with a heatsink and cooling fans. **c)** Image of assembled 24-well LAVA board, with optical, cooling, and electronics subsystems highlighted.

### Optimization and characterization of illumination uniformity

We established the LAVA optical system design by modeling the optical configuration of a single well in the optical ray tracing software Zemax OpticStudio (Figure S2A-S2H). We then optimized the optical configuration for uniform well illumination (Figure 2A). Due to spatial constraints of a TC incubator, we used optical diffusers, rather than lenses, in addition to optical scattering from the 3D-printed light guides to ensure illumination uniformity. In the Zemax model, parameters such as LED position on the circuit board, diffuser strength, and light guide dimensions were optimized to reduce intensity drop-off at the well edge (Figure S2A-S2H, Figure S3A-S3E). Modeling results showed that the parameter with the strongest effect on uniformity was the axial thickness, *d*, of the two 3D-printed light guides (labelled in Figure 1B). Based on these modeling results, we fabricated LAVA devices and experimentally validated the resulting well uniformity by imaging LAVA wells under a low-magnification microscope (Figure 2B, Figure S4A-S4C). Measurement of light intensity as a function of radial distance confirmed the improved illumination uniformity, which came at the expense of maximum illumination intensity (Figure 2B). Increasing *d* from 1 cm to 1.5 cm attenuated the intensity decrease at the well edge from 20.4% to 16.9%, i.e. a roughly 20% improvement in uniformity. A larger *d* also resulted in a two-fold improvement in well-to-well variability between the 24 independent wells (2.6% versus 1.2% coefficient of variation) (Figure S4D). For experimental applications where intensity of illumination is paramount to well uniformity, a lower *d* could be used to achieve higher illumination intensities. Thus, the LAVA boards can be used in two hardware configurations, which are summarized as follows: (1) a low-intensity, high-precision configuration at *d* = 1.5 cm where intensity can be programmed from 0 – 10µW/mm^2^ in 0.0024 µW/mm^2^ increments with high illumination uniformity and low well-to-well variability and (2) a high-intensity, low precision configuration at *d* = 1 cm where well intensity resolution, variability, and uniformity are sacrificed to achieve a doubling in light intensity (0 – 20µW/mm^2^ in 0.005 µW/mm^2^ increments) (Figure 2C).

**Figure 2.**
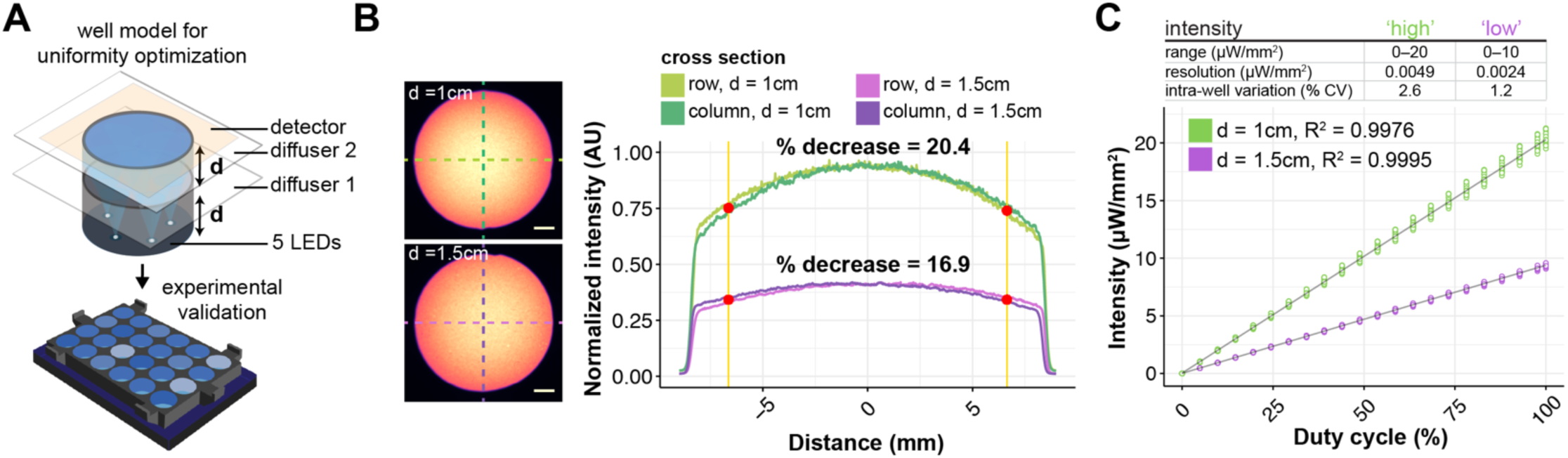
Optical design for illumination uniformity of TC plate wells. **a)** Schematic of Zemax model used for LAVA optical system optimization. **b)** Brightfield images of LAVA board wells (left) and graph of intensity linescans along indicated cross-sections (right) characterize the intensity uniformity of the 24-well LAVA device under two configurations, where light guide thickness, *d*, is either (1) 1 cm (top, green) or (2) 1.5 cm (bottom, purple). Percent decrease is calculated between intensity at center of well and intensity at highlighted red point, which indicates location of well edge of a 24-well culture plate. Graph shows mean normalized intensity over 4 independent wells. Scale bar 2.5 mm. **c)** Measured light intensity in response to the programmed duty cycle of the LED pulse-width modulation signal. Graph shows measured intensity from each well of a 24-well LAVA board and curve fit to a linear regression model.

## RESULTS

### Intensity control of optogenetic stimulation reveals Brachyury expression level is dependent on LRP6 oligomer number and size

During mammalian embryonic development, gradients in Wnt signal intensity control the progression of cell lineage commitment and axis patterning (Zeng et al. 1997; Liu et al. 1999; Kimura-Yoshida et al. 2005; Arnold & Robertson 2009). In hESCs, the strength of Wnt signaling similarly modulates cell lineage commitment and differentiation potential (K. C. Davidson et al. 2012; Blauwkamp et al. 2012; Sumi et al. 2008; Bernardo et al. 2011). Equipped with a method for optogenetic stimulation of cell cultures, we used the LAVA boards to activate canonical Wnt/β-catenin signaling in a clonal hESC line expressing the optoWnt system (Repina et al. 2019). In addition to on/off control of Wnt signaling, we sought to determine whether optoWnt could be activated in a dose-responsive manner to better mimic the signal gradients present during development.

Since Cry2 oligomerization is a dynamic, reversible process wherein clustering is triggered upon photon absorption (Duan et al. 2017; Bugaj et al. 2013), we reasoned that the fraction of photostimulated Cry2 proteins per cell could be controlled with light intensity (Figure 3A). Using the LAVA boards, we were able to set independent wells to different light intensities using pulse width modulation (PWM) (Figure 2C). Upon continuous photostimulation at variable intensities, we indeed observed that the number and size of visible LRP6 oligomers per cell increased monotonically with illumination intensity (Figure 3B, Figure S5A-S5B). To determine whether increased Cry2 oligomerization translated to a stronger Wnt signal intensity, we probed for expression Brachyury (BRA, also known by its gene name, *T*), a direct transcriptional target of Wnt signaling and regulator of mesendoderm and primitive streak differentiation (Rivera-Pérez & Magnuson 2005; Yamaguchi et al. 1999; Arnold et al. 2000; Lindsley et al. 2006). Following a similar trend to Cry2 oligomerization, the mean intensity of BRA immunostaining increased with light intensity, suggesting that an increased number and size of LRP6 clusters results in a stronger differentiation signal (Figure 3C, Figure S5C).

**Figure 3.**
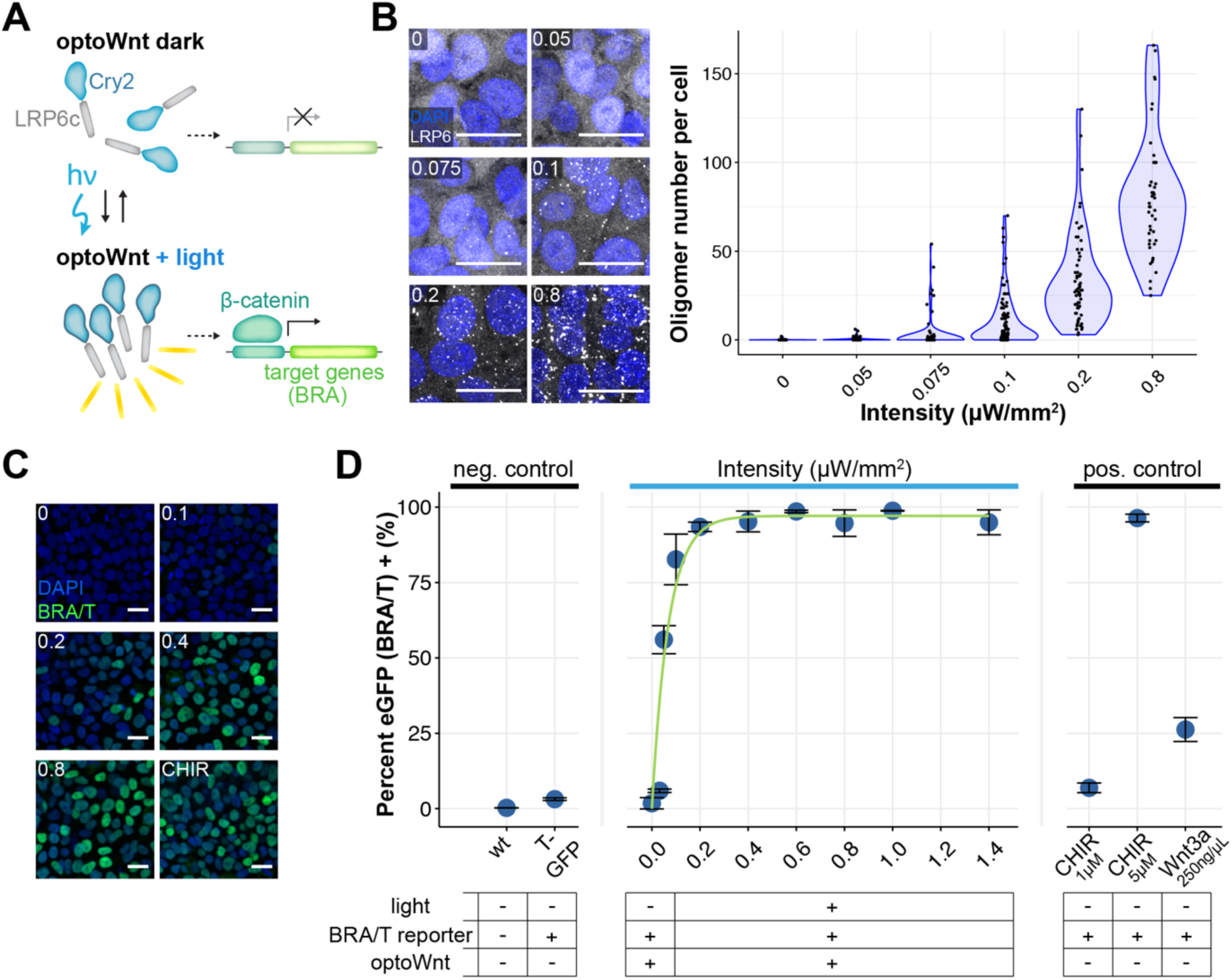
Optogenetic induction of BRA expression is light dose dependent. **a)** Schematic of optoWnt system. In the dark, the Cry2 photosensory domain is diffuse. Illumination induces LRP6c oligomerization and transcription of β-catenin target genes. **b)** Immunostaining for LRP6 (left) and quantification of cluster number per hESC in response to increasing light intensity after 1 hr illumination. Graph shows individual cell quantification, sith each point representing a single cell. Scale bar 25 µm. **c)** Immunostaining for BRA in response to increasing light intensity after 24 hr illumination or 3 µM CHIR treatment. Scale bar 25 µm. **d)** Flow cytometry of optoWnt hESCs expressing eGFP reporter for BRA/T treated with varying light intensities or with Wnt pathway agonists (Wnt3a recombinant protein or CHIR). Graph shows percent eGFP positive cells and nonlinear least squares fit to increasing exponential decay curve. Subset of data reproduced from Repina et al. 2019. Graph shows mean ± 1 s.d., n = 3 biological replicates.

To better quantify BRA expression at a single-cell level, we used a clonal hESC cell line co-expressing the optoWnt system and an eGFP reporter driven by the endogenous *T* locus (Repina et al. 2019). Live-cell analysis with flow cytometry showed a heterogeneous response to Wnt stimulation at both low (0.1 µWmm^-2^) and high (1.4 µWmm^-2^) light intensities, confirming BRA immunostaining results (Figure S5C-S5D). Maximal light-induced activation increased BRA expression by ∼33-fold over unilluminated optoWnt hESCs, which notably showed no detectable activation in the dark. Quantification of the percentage eGFP-positive cells exhibited an exponential increase with light intensity that greatly exceeded activation achieved with recombinant Wnt3a protein (Figure 3D). Fitting to an increasing exponential decay function showed a very rapid increase in BRA expression at low light intensities (time constant τ = 0.07 µWmm^-2^ / percent eGFP+, see methods), with saturation reached at ∼0.4 µWmm^-2^. The high sensitivity at lower light intensities suggests a binary switch for onset of BRA expression above a signaling threshold, followed by a monotonic increase in BRA expression levels in a light dose-dependent manner.

### Analysis of phototoxicity reveals no detectable effects on hESC viability below a 1 µWmm^-2^ illumination intensity threshold

Since high intensity light can induce phototoxicity in cells (Tyssowski & Gray 2019; Acker et al. 2016; Allen et al. 2015; Yizhar et al. 2011; Owen et al. 2019), we used the LAVA boards to analyze the phototoxicity threshold for hESC cultures. Inefficiencies in LED semiconductors cause heating, and though we incorporated a heat sink, cooling fans, and electrical vias in the circuit board design to conduct heat away from the cell culture plate, we observed heating of the LED dyes at continuous operation at high intensities (Figure S6A). Within the intensity range where we did not observe significant heating (0 – 2 µWmm^-2^), we assessed potential phototoxic effects on hESC cultures after 48 hrs of continuous illumination. Above 1 µWmm^-2^, we observed a significant increase in apoptosis and membrane integrity markers Annexin IV and propidium iodide (PI), as well as a corresponding increase in cell debris and edge roughness of hESC colonies (Figure S6B-S6C). Thus, for all optogenetic stimulation we used light intensity under 1 µWmm^-2^ (specifically, 0.8 µWmm^-2^). At this intensity, we saw no decrease in pluripotency markers SOX2, NANOG, and OCT4 after 48 hrs of illumination of wild-type hESCs, as well as no spontaneous BRA+ mesendoderm differentiation (Figure S6D). Notably, since saturation in the percentage BRA+ optoWnt cells occurred at ∼0.4 µWmm^-2^ (Figure 3D), the optoWnt operating range falls below the 1 µWmm^-2^ phototoxicity threshold.

### Temporal control of optoWnt shows BRA downregulation after light withdrawal

In addition to intensity control, LAVA boards enable temporal control of illumination patterns (Figure 4A). Given the reversible oligomerization of Cry2 (Bugaj et al. 2013), we reasoned that the optoWnt system can be readily applied to regulation of Wnt temporal dynamics.

**Figure 4.**
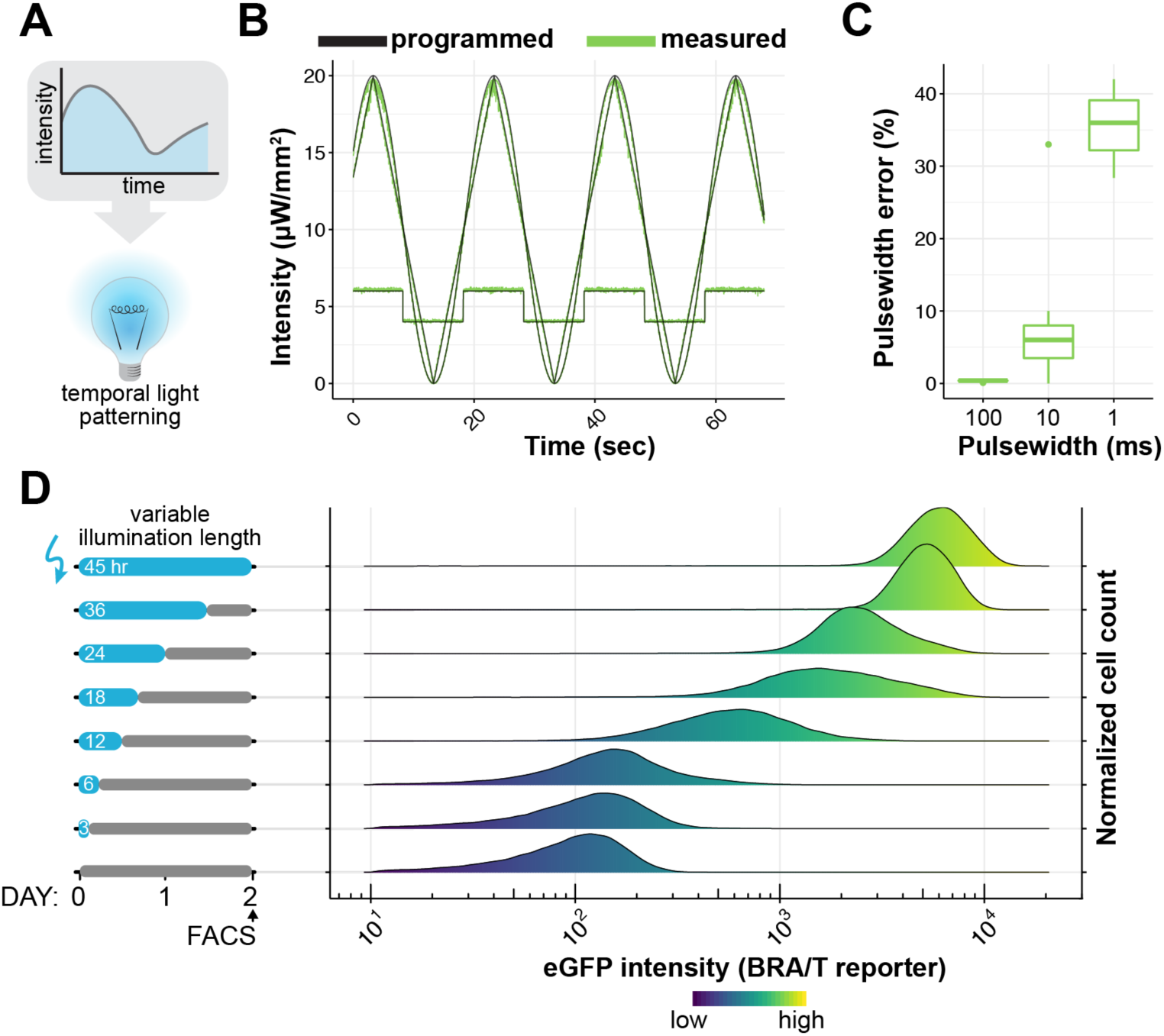
Characterization of temporal control using LAVA devices. **a)** Schematic of temporal light patterning in optogenetics. **b)** LAVA board well intensity as a function of time of various waveforms programmed through LAVA GUI. Programmed values shown in black, measured intensities shown in green. **c)** Percent error in measured pulsewidth relative to programmed pulsewidth for blinking sequences programmed through the LAVA GUI at 50% duty cycle and at indicated pulsewidth on-times. **d)** OptoWnt hESCs containing an eGFP reporter for endogenous BRA/T activity were illuminated for varying lengths of time and analyzed by flow cytometry at a fixed endpoint. Graph shows histograms of eGFP expression at each illumination condition. Cell count histograms normalized to total cells per condition (∼30,000 cells), n = 3 biological replicates.

We designed a LAVA board GUI to allow users to input the desired temporal light patterns for each independent well (Figure S7A-S7C). After input of illumination parameters, the GUI wirelessly transmits the parameters to the LAVA device in the form of a JSON file. The file is parsed by the on-board Raspberry Pi microcontroller, which then to sets LED intensities and temporal patterns for the duration of the experiment. The user can set each of the 24 wells to one of three modes: (1) constant illumination at a specified intensity; (2) blinking at a specified intensity, duty cycle, and period; and (3) a series of linear or sinusoidal functions at specified function parameters. Multiple piecewise functions can be programmed in sequence, enabling an immense variety of complex, time-varying light patterns (Figure 4B, **Video S1**). The shortest possible blink, i.e. the temporal resolution of the device, is set to 1 ms in firmware, though we observed a significant drop in the accuracy and precision of stimulation at pulsewidths below 10 ms (Figure 4C, Figure S8).

We next asked whether sustained optoWnt activation is required to maintain BRA expression during hESC differentiation. We thus illuminated optoWnt cells expressing the T/eGFP reporter with varying durations of light and quantified eGFP fluorescence with flow cytometry at a fixed endpoint (Figure 4D). The duration of optoWnt stimulation determined endpoint eGFP intensity, suggesting that sustained illumination and optoWnt activation is necessary for a sustained β-catenin transcriptional response.

### Spatial localization of Wnt signaling and hESC differentiation as a model for early embryonic Wnt patterning

Another advantage of optogenetic control is the ability to manipulate the spatial location of signal activation (Figure 5A). Though precise patterned illumination can be achieved with confocal scanning or the use of spatial light modulators (Repina et al. 2017), the cost and complexity of such microscope systems are restrictive, particularly for longer biological processes and/or large sample sizes. Instead, the majority of optogenetic studies leverage genetic specificity to target cell types of interest (Yizhar et al. 2011) or subcellular tagging to specific cell compartments (Benedetti et al. 2018) to achieve higher spatial resolutions. To easily incorporate spatial light patterning during cell culture, we designed die-cut photomasks that can be adhered to TC plates during illumination (Figure S9A). The mask feature size was limited by the cutting resolution of the die cutter to ∼150 µm (Figure 9B). Using such photomasks, we were able to illuminate wells with arbitrary light patterns and induce optoWnt clustering only in illuminated regions (Figure 5B). Since the mask was placed underneath the TC plate, we anticipated that there would be light scattering through the plate bottom (170 µm-thick coverglass) that would compromise the mask resolution (Figure S9C). To quantify the extent of light scattering, we measured the number of LRP6 clusters as a function of distance beyond the mask edge and found that clusters were induced ∼50 µm from the mask edge (Figure 5C). Thus, photomask feature size was limited to ∼150 µm while patterning resolution (full width at half maximum, see methods) was ∼100 µm (Figure 5C).

**Figure 5.**
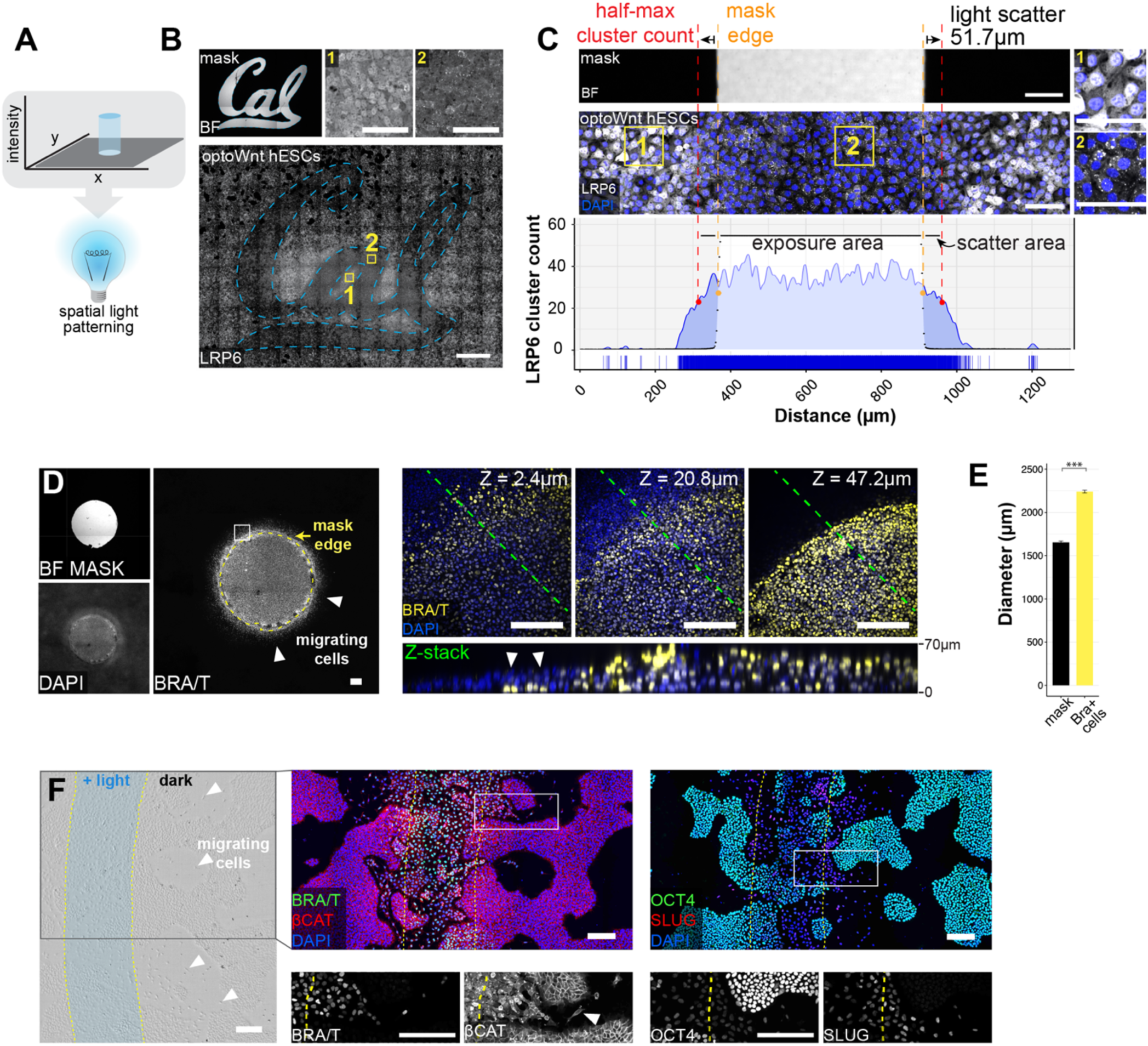
Spatial light patterning with LAVA devices for localized optoWnt activation **a)** Schematic of spatial light patterning in optogenetic experiments. **b)** Stitched brightfield and fluorescence confocal images of optoWnt hESCs illuminated with UC Berkeley (Cal) logo photomask. Immunostaining for LRP6 oligomers, with representative image of masked region (1) and illuminated region (2) as shown. Scale bar 100 µm (top), 1 mm (bottom). **c)** Quantification of light scattering through bottom of TC plate shows a ∼50 µm spread (full width at half max, red line) of optoWnt oligomers beyond photomask edge (orange line). Figure shows brightfield image of photomask (top), fluorescence image of immunostaining for LRP6 (middle), and quantification of LRP6 cluster count (bottom). Insets (1) and (2) show masked and illuminated regions, respectively. Scale bars 100 µm. **d)** Patterned illumination with a 1.5 mm diameter circle of light. BRA immunostaining with photomask overlay shown in left panel. Confocal z-stacks of bottom (closest to coverslip, Z = 2.4 µm), middle (Z = 20.8 µm) and top (Z = 47.2 µm) cell layers show BRA+ cells localized beyond photomask boundary and under the epithelial cell layer (white arrows). Bottom panel shows z-slice through cross-section highlighted with green line. Scale bars 100 µm. **e)** Quantification of BRA+ cell localization beyond photomask edge. Mean diameter of circular photomask pattern was quantified using the brightfield image channel, and mean diameter of BRA+ cell pattern was quantified through immunostaining, as shown in (d). Graph shows mean measured diameter ± 1 s.d., n = 3 biological replicates. Student’s t-test (two-tail). Scale bars 200 µm. **f)** Patterned illumination with stripe of light, 500 µm in width. Brightfield image (left panel) with overlay of light pattern shows cells with mesenchymal morphology localized beyond of region of illumination (white arrows). Immunostaining for BRA and β-CAT (middle panel); OCT4 and SLUG (right panel). Overlay of light pattern is highlighted in yellow, and zoom-in (white box) shown below. Scale bars 200 µm.

During mammalian development, a gradient of Wnt signaling emerges across the epiblast (Arnold & Robertson 2009; Liu et al. 1999; Rivera-Pérez & Magnuson 2005). Subsequently, cells in the posterior epiblast break away from the epithelial cell layer, migrating as single cells, which results in spatial rearrangement and morphological symmetry-breaking of the epiblast in the region of the primitive streak (Thiery & Sleeman 2006; Thiery et al. 2009; Williams et al. 2012). We modeled this spatial pattern of Wnt signaling by activating optoWnt hESCs in defined regions within the culture. To determine whether optoWnt activation could trigger cell migration outside the area of illumination, we used the LAVA boards to illuminate cells with a 1.5 mm-diameter circular light pattern (Figure 5D). Indeed, immunostaining showed that BRA+ cells localized both within the illuminated region as well as up to ∼300 µm beyond the illumination boundary (Figure 5E). Given the ∼50 µm photomask scattering (Figure 5C), we excluded the possibility that cells were activated by scattered light. Intriguingly, within the illuminated circle, BRA+ cells grew vertically upwards and piled four to six cell diameters in height (∼70 µm). In contrast, BRA+ cells beyond the illumination boundary localized underneath the epithelial cell layer (Figure 5D). These cells likely underwent an epithelial-to-mesenchymal transition (EMT) and migrated below the hESC colony, reflecting a migratory phenotype that has been analogously observed in hESCs undergoing EMT in conditions of high cell density and confinement (Warmflash et al. 2014).

We next illuminated optoWnt cultures with a stripe of light to mimic the spatial geometry of Wnt gradients established across the epiblast (Rivera-Pérez & Magnuson 2005). Cells in the illuminated stripe broke away from hESC colonies, adopted a mesenchymal morphology, and localized up to 500 µm beyond the boundary of the light pattern (Figure 5F). Immunostaining confirmed that these migratory cells expressed BRA, while surrounding unilluminated cells retained epithelial morphology with no detectable BRA expression. In addition, migratory cells showed a decrease in OCT4 expression, a shift in β-CAT localization away from the plasma membrane, and an increase in SLUG expression, all consistent with cells undergoing an EMT (Figure 5F). Taken together, these data show that optogenetic Wnt activation is sufficient for inducing a migratory cell phenotype and that patterned illumination can be used as a tool to further study Wnt patterning and gastrulation-like events in ESC culture.

## DISCUSSION

Spatial and temporal morphogen gradients have long been known to exist during development, though there has been a paucity of tools to create and perturb such signaling patterns. In this study, we combined optogenetic control of Wnt signaling with novel engineered illumination devices, LAVA boards, to dynamically control hESC morphogen signaling in intensity, space, and time. Using this precise manipulation of morphogen signaling, we show that optogenetic Wnt patterning can be used to mimic embryonic morphogen patterning *in vitro*. This platform will enable future studies that elucidate spatial and temporal Wnt signaling thresholds and mechanisms of cell patterning in a variety of biological systems.

### LAVA boards for quantitative, spatiotemporal control of Wnt signaling dynamics

With the continued development of new optogenetic proteins that perturb cell signaling and protein-protein interactions, there is a need for illumination devices that allow high-throughput intensity, spatial, and temporal control of stimulation. Previous illumination technologies for multi-well plate photostimulation have helped advance optogenetic studies, but have addressed only some of these individual design criteria and do not fully characterize intensity parameters, uniformity, spatial and temporal resolution, or undesired heating and toxicity (Gerhardt et al. 2016; E. A. Davidson et al. 2013; Lee et al. 2013; Hannanta-Anan & Chow 2016; Bugaj et al. 2018; Hennemann et al. 2018). Thus, in designing LAVA boards, we addressed such concerns and engineered devices that programmatically control photostimulation intensity, timing, and location at 0.005 µWmm^-2^, 10 ms, and 100 µm resolution, respectively. As optical system cost and ease of use can be significant barriers for adoption of optogenetic studies, we also developed a GUI for simple configuration and wireless upload of desired intensity patterns to LAVA devices from a personal computer. Further, to make optogenetic studies more accessible, we provide a detailed protocol, design files, and software source code for both 24-well and 96-well plate LAVA board assembly. Each device takes ∼8 hrs and less than $500 to fabricate and assemble.

One of the major optical design parameters we optimized during LAVA board design was illumination uniformity. Since the strength of induced Wnt signaling is dependent on light dosage (Figure 3D), nonuniformity can give rise to signal variation across the region of optogenetic stimulation. Specifically, the optoWnt dose response showed that Wnt activation can be sensitive to up to 0.025 µWmm^-2^ differences in intensity, especially at photostimulation near 0.1 µWmm^-2^ (Figure 3D). Despite this high sensitivity, the optimized LAVA board design is equipped to stimulate wells at sufficiently uniform levels. Illumination at 0.1 µWmm^-2^, for example, would have a 0.017 µWmm^-2^ intensity drop-off at the well edge (16.9% decrease) (Figure 2B), which is below the 0.025 µWmm^-2^ design criterium and thus results in uniform Wnt signal activation within the well. Though characterization of uniformity is critical for controlled signal activation, surprisingly this design parameter has not been addressed in previous illumination systems (Gerhardt et al. 2016; Olson et al. 2014).

Equally important to illumination uniformity is programmable control of intensity for dose-responsive and time-varying signal activation. Because we observed dose-dependent Wnt signal activation in response to increasing light intensities (Figure 3D), the LAVA boards combined with the optoWnt system can be readily applied to various biological systems where Wnt signal strength directly regulates cell fate outcome. For example, different Wnt signal intensities can drive dose-dependent or divergent lineage specification during neural fate commitment (Kirkeby et al. 2012; Kiecker & Niehrs 2001), hematopoiesis (Luis et al. 2011), and mesoderm differentiation (K. C. Davidson et al. 2012; Kempf et al. 2016). In addition, the optoWnt dose response suggests a role for LRP6 oligomer size in regulating β-catenin degradation (Kim et al. 2013; Li et al. 2012), which can be used to further shed light on the molecular regulation of Wnt signaling. In parallel, temporal modulation with LAVA devices opens a range of possible studies of Wnt signal dynamics (Figure 4). Combined with optoWnt reversibility (Bugaj et al. 2013)(Figure 4D), intricate studies of Wnt signaling thresholds and timing of signaling oscillations during development (Bao et al. 2016; Aulehla et al. 2003; Lian et al. 2012; Yu et al. 2008) can now be performed with much greater ease compared with microfluidic approaches (Sonnen et al. 2018) or manual pipetting (Massey et al. 2019).

Lastly, for spatial control, we implement a dye-cut photomask to achieve spatial patterning at 100 µm resolution (Figure 5C), an improvement relative to previous innovative photomask approaches (Müller et al. 2014; Shao et al. 2018). Patterned illumination thus enables the unprecedented ability to ‘paint’ Wnt activation onto target cells of interest and study how the shape, size, and intensity of spatial patterns influences differentiation and morphogenesis. Elegant optogenetic approaches for studying spatial signaling thresholds, like patterned ERK signaling during early *Drosophila* embryogenesis (Johnson & Toettcher 2019; Johnson et al. 2017), can now be extended to morphogen patterning in stem cell-based and organoid models for development.

### Limitations

Having established proof of principle applications for LAVA devices and the optoWnt system, we would like to comment on their limitations as well as considerations for generalizability to other optogenetic systems. Many hardware components of LAVA boards are exchangeable, so that LEDs of other colors can readily be incorporated to activate red or UV-based light sensors. Currently, the hardware design allows independent control of a single color, but can be extended to multicolor stimulation by modifying the electronics design to incorporate more LED drivers.

To stimulate photosensory proteins, light delivered by LAVA devices must be of sufficiently high intensities. We have found that the achieved illumination intensities (0 – 20 µWmm^-2^) are sufficient for optoWnt photostimulation, which requires merely 0.5 – 1 µWmm^-2^ for maximal activation (Figure 3D). However, LAVA board intensities may be insufficient for less sensitive photosensory domains that can require pulses of up to 1,000 – 10,000 µWmm^-2^ (Yizhar et al. 2011). LAVA board intensities can be increased by minimizing the light guide thickness, *d* (see Figure 2), but the maximal illumination intensity is still limited by the maximum forward current of LED dyes, as well as by the phototoxicity of light at higher intensities (Tyssowski & Gray 2019; Acker et al. 2016; Allen et al. 2015; Yizhar et al. 2011; Owen et al. 2019). Thus, for high-intensity applications, microscope-based systems and laser illumination would be more suitable.

Lastly, spatial patterning with a photomask is limited to a static, two-dimensional light pattern, with resolution limited by light scattering through the TC plate bottom surface. In the future, the pattern feature size can be improved by using chrome masks or laser-plotted mylar masks, which are limited to sub-micron and ∼5 – 10 µm features, respectively (Folch 2016). Such methods can also be used to achieve greyscale modulation (Folch 2016). In summary, LAVA devices are versatile tools for photostimulation of cell cultures, and can be applied to many biological systems for studies of cell signaling dynamics, high-throughput screens, and control of protein-protein interactions.

## ACKNOWLEDGEMENTS

We thank Laura Waller for discussions on optical design, as well as Nicholas Antipa and Zachary Phillips for helpful discussions on Zemax modeling and uniformity characterization. We are grateful to Christopher Myers, Mitchell Karchemsky, and Rundong Tian at the UC Berkeley CITRIS Invention Lab for assistance and discussions of rapid prototyping design and techniques. We also thank members of the D.V.S lab for helpful discussions. We are grateful to Mary West from the QB3 High-Throughput Screening Facility and CIRM/QB3 Shared Stem Cell Facility and Hector Nolla from the CRL Flow Cytometry Facility for technical assistance. Funding supporting this work was provided by the US National Institutes of Health (R01NS087253 to D.V.S.), the U.S. National Science Foundation (to N.A.R.), the Siebel Scholars Foundation (to N.A.R.).

## AUTHOR CONTRIBUTIONS

N.A.R. conceived the study, designed and performed experiments, performed analysis, and wrote the manuscript. T.M. designed software control and graphical user interface. X.B. performed experiments. R.S.K. conceived the study. D.V.S. conceived the study and wrote the manuscript.

## DECRALATION OF INTERESTS

N.A.R., T.M., and D.V.S are co-inventors on related intellectual property.

## METHODS

### Zemax modeling and uniformity optimization

The ray-tracing software Zemax OpticStudio was used in Non-Sequential mode to model illumination of a 24-well plate. Based on modeling results, the optimized configuration parameters are as follows: 5 surface-mount LEDs are symmetrically radially distributed around a 5mm-radius circle; the radius of each light guide is 8.25 mm; one 80° circular optical diffuser is placed between the two light guides and another onto the top light guide (i.e. between light guide and TC plate); the thickness of each light guide is 1.5 cm; and light guides are manufactured from black 3D-printed acrylic.

### LAVA device construction

LAVA devices are constructed using two custom printed circuit boards (PCB) designed in EAGLE (Autodesk). PCB1 contains electronics for LED control while PCB2 is the power distribution board. For 24-well illumination, PCB1 contains solder pads for a circular array of 5 LEDs per well, which are connected in series and illuminate each well through two 3D-printed light guides and a series of diffusers (optical configuration optimized in Zemax, see below). For 96-well illumination, PCB1 contains solder pads for 1 LED per well of a 96-well plate; given the 24-channel LED driver, independent illumination control is possible for each group of 4 wells. For each channel, the ground wire connects to TLC5947 driver and is modulated with pulse-width modulation, while the positive terminal connects to the power plane of PCB1. PCB1 also contains headers for electrical connection to cooling fans. A heatsink mounts onto the bottom of PCB1, using thermally conductive adhesive (Arctic Silver, ASTA-7G), in a region without silk screen and thermally conductive electrical vias that draw heat away from surface-mount LEDs.

In PCB2, a power supply connects through a barrel power jack to power the LEDs through an LED driver (TLC5947, Adafruit). Power is also supplied to three fans and the Raspberry Pi microcontroller through switching voltage regulators.

On top of PCB1, optical assembly and TC plate is mounted in such a way that TC plate is illuminated from the bottom. It is critical that TC plate is made of black, opaque plastic with a thin, 170 µm coverslip bottom (Eppendorf Cell Imaging Plate, 24-well) to avoid light bleed-through between wells and thus enhance high spatial patterning resolution. The LED driver, PCB2, and the Raspberry Pi microcontroller are all mounted and electrically connected to PCB1, and the entire assembly is mounted onto an acrylic laser-cut base through vibration-dampening mounts. The base contains rubber footpegs to reduce static or electrical shorting with the TC incubator racks.

For details of LAVA board design and fabrication, see supplemental methods which include a detailed protocol, design files, and software source code for 24-well and 96-well plate LAVA board assembly. In brief, LAVA boards were constructed in the CITRIS Invention Lab, a UC Berkeley rapid prototyping facility, using the following equipment: 3D printer (Ultimaker 3), laser cutter (Speedy 400, Trotec), and standard soldering tools. Photomasks were dye-cut on a vinyl cutter (CAMM-1 GS-24, Roland).

### LAVA software control and graphical user interface

The LEDs are controlled by an Adafruit 24-Channel 12-bit PWM LED driver with an SLI interface to a Raspberry Pi Zero W. The 12-bit PWM resolution allows 4086 unique illumination intensity levels over the LED operating range. For ease of use, a GUI has been written in Java and is conveniently packed into an executable file. This interface allows for independent control of each of the 24 channels. To accommodate the variety of experimental conditions, each LED can be programmed to a constant illumination, a blinking pattern, or a series of linear and sinusoidal patterns. Since each board has slightly different intensity characteristics, the intensity to PWM calibration parameters are input at runtime. Sinusoidal and linear functions are interpolated at a frequency of 1 Hz whereas blinking patterns have been tested up to 100 Hz. Since the LED board’s USB port may be inaccessible during certain experiments, it is possible to wirelessly upload new illumination settings from any Wi-Fi capable computer. The Java program parses the illumination settings and PWM calibration parameters, packages them into a JSON file, and transmits these settings to the Pi over an SFTP channel.

Upon booting the Raspberry Pi, a C++ script executes, checks the device for previous illumination settings and resumes the patterned illumination if found. The Pi polls for changes in the JSON file every few seconds, so the changes of a newly uploaded pattern will be reflected without an additional reboot. It should be noted that the decision to use C++ was motivated by a desire to break through certain speed limitations posed by an interpreted language’s rate of execution. A Python implementation was completed and included on the Pi operating system to easily generate custom pattern scripts, but only the compiled C++ version is able to drive the 24 channels at the desired refresh rate.

### LAVA device intensity, uniformity, heating, and spectral characterization

For LAVA board characterization, light intensity was measured with a power meter (PM100D, Thorlabs) with a photodiode power sensor (S121C, Thorlabs) at 470 nm. For high-resolution measurement of temporal light patterns, the Thorlabs power meter was connected to an oscilloscope. Well uniformity measurements were performed by imaging wells on a Nikon Z100 microscope with a wide-field lens (Nicole AZ-Plan Apo 1x) and sCMOS monochrome camera (pco.edge 5.5). Image intensity was quantified as a function of distance in Fiji(Schindelin et al. 2012). Well temperature was measured with a digital multimeter (87-V, Fluke) with a Type-K thermocouple probe (80BK-A, Fluke). LED emission spectra were measured with a spectrometer (Red Tide USB650, Ocean Optics).

### Embryonic stem cell culture

For routine culture and maintenance, all optogenetic and wild-type hESCs (H9, WiCell) (Thomson et al. 1998) lines were grown on Matrigel (Corning, lot # 7268012, 7275006) coated plates in mTeSR1 medium (STEMCELL Technologies) and 1% penicillin/streptomycin (Life Technologies) at 37°C and 5% CO_2_ with daily media changes. Optogenetic cells were cultured with hood lights off. For illumination experiments, cells were singularized with Accutase (STEMCELL Technologies) at 37°C for 5min and seeded onto Matrigel-coated 24-well plates (0030741021, Eppendorf, black-walled with 170 µm coverglass bottom) in E8 media (STEMCELL Technologies) with 5 µM ROCK inhibitor Y-27632 (Selleckchem) at a density of 35k – 70k cell cm^-2^. Wnt agonist CHIR99021 (Stemgent) or Wnt3a protein (StemRD) was diluted in E8 media and added to cells. Clonal optoWnt knock-in lines and clonal *BRA/T* reporter lines were generated through CRISPR/Cas9-mediated recombination as previously described (Repina et al. 2019).

### Immunostaining and imaging

Cells were fixed with 3% PBS – paraformaldehyde (ThermoFisher) for 20min at room temperature and subsequently washed three times with PBS. Blocking and permeabilization was done with 5% donkey serum (D9663, Sigma-Aldrich) and 0.3% Triton X-100 (Fisher Scientific) (PBS-DT) for 1 hour. Cells were incubated with primary antibodies (Supplementary Table 1) at 4°C overnight, then washed three times with PBS, and incubated with fluorescently conjugated secondary antibodies (Invitrogen) at 1:250 dilution for 1 hour at room temperature. Both primary and secondary antibodies were diluted in PBS-DT. Cells were washed with PBS and stained with 0.1 µg mL ^-1^ DAPI nuclear stain (ThermoFisher) prior to imaging. Confocal imaging was performed on a Perkin Elmer Opera Phenix system (QB3 High-Throughput Screening Facility). Brightfield and widefield fluorescence imaging was performed on a Zeiss AxioObserver epi-fluorescent microscope (CIRM/QB3 Shared Stem Cell Facility).

### Image analysis

Microscopy image processing, including stitching and z-slice projection, was performed in Fiji (Schindelin et al. 2012) and image quantification was performed in CellProfiler (Carpenter et al. 2006) with custom analysis pipelines detailed below.

For quantification of BRA, SOX2, OCT4, and NANOG nuclear fluorescence intensity, nuclei stained with DAPI were identified and used to generate a binary mask applied to the appropriate fluorescence channel. The mean fluorescence intensity per cell nucleus was calculated for each cell in a given field of view. A threshold defining ‘positive’ cells was determined from signal intensity of negative control wells. Quantification of LRP6 oligomer size was similarly performed in CellProfiler using the IdentifyPrimaryObjects module.

Spatial light patterning resolution was quantified by illuminating optoWnt hESC wells with a dye-cut photomask for 1 hr at 1 µWmm^-2^ and immunostaining for LRP6. With mask attached to plate, wells were imaged in brightfield at varying focus positions, to capture images of both the photomask and cells. The mask was subsequently removed, and cells were imaged in brightfield and fluorescence channels (with 5x and 60x water-immersion objectives on a Perkin Elmer Opera Phenix confocal system). The mask image was aligned to the LRP6 fluorescence channel using the brightfield channel in Fiji (Schindelin et al. 2012). An identical alignment procedure was used for all spatial patterning data shown. For quantification of light scattering, multiple fields of view were stitched together and oligomers were manually counted in Fiji based on the LRP6 fluorescence channel. The pixel location of each identified LRP6 oligomer was recorded. Since the photomask pattern was a vertical stripe, the oligomer counts were summed vertically over multiple fields of view to quantify the spatial distribution of oligomers relative to the photomask. The spatial extent of photostimulation due to light scatter was calculated by determining the full width half maximum (FWHM) of the LRP6 oligomer distribution outside of the photomask.

### Flow cytometry and analysis

Cells were lifted with Accutase at 37°C for 5min, centrifuged, and resuspended in flow buffer (0.5% bovine serum albumin in PBS (w/v)) for analysis. For photoxicity experiments, centrifuged cells were resuspended in PBS and stained with AlexaFluor 488-conjugated annexin V and 1 µg mL^-1^ propidium iodide (PI) as per kit manufacturer recommendations (V13241, ThermoFisher). Cells were analyzed on a BD LSR Fortessa X20 (UC Berkeley Cancer Research Laboratory Flow Cytometry Facility). Cell counting for proliferation assays was performed using a ThermoFisher Attune (UC Berkeley CIRM/QB3 Shared Stem Cell Facility) by measuring the number of cells per unit volume.

Data analysis was performed with FlowJo 10 software. To determine the fraction of BRA-eGFP+ cells after light treatment, gating was set such that less than 0.5% of undifferentiated wild-type hESCs were positive for eGFP and mCherry. Data were fit to an increasing exponential decay curve using a nonlinear least squares model in R:

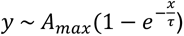

where *y* represents the percent BRA-eGFP+ cells and *x* is light intensity (µWmm^-2^). The curve asymptotically approaches the maximum percentage of BRA-eGFP+ cells (*A_max_* = 97.1%) with time constant τ = 0.07.

### Statistical analysis and graphing

Data are presented as mean ± 1 standard deviation (s.d.) unless otherwise specified. Statistical significance was determined by Student’s t-test (two-tail) between two groups, and three or more groups were analyzed by one-way analysis of variance (ANOVA) followed by Tukey test. P < 0.05 was considered statistically significant (NS P>0.05, *P<0.05, **P<0.01, ***P<0.001).

## SUPPLEMENTARY DATA

This section includes:

**Supplementary Table 1.**
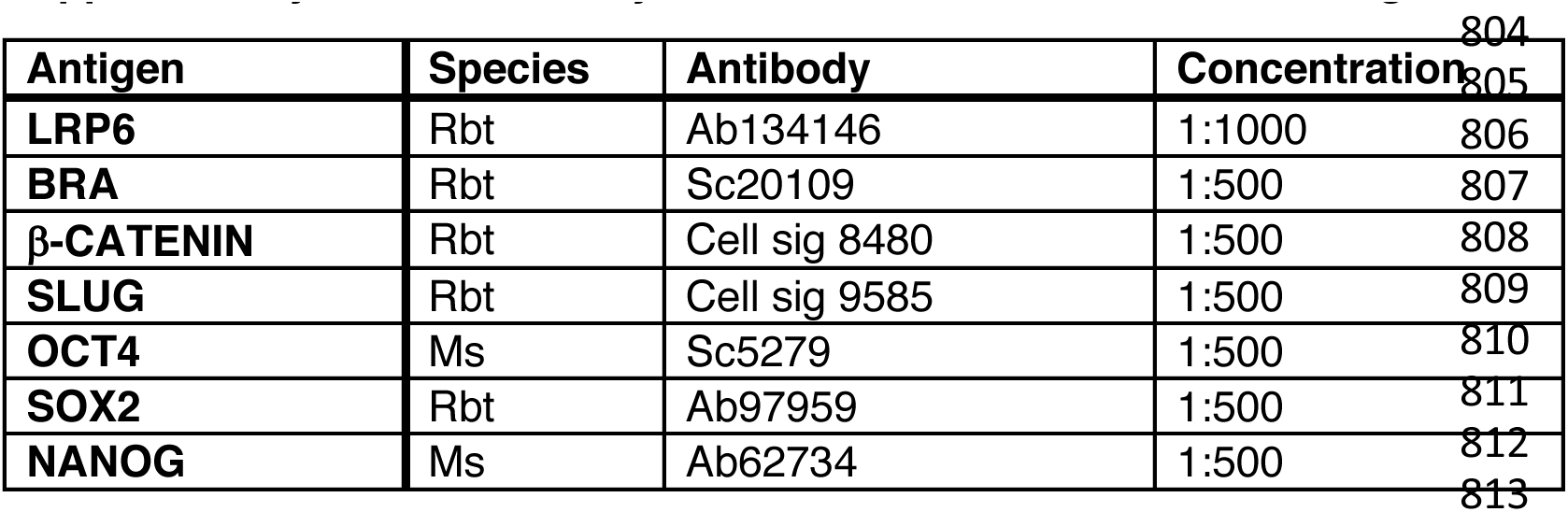
Primary antibodies used for immunostaining.

| Antigen | Species | Antibody | Concentration |
| --- | --- | --- | --- |
| LRP6 | Rbt | Ab134146 | 1:1000 |
| BRA | Rbt | Sc20109 | 1:500 |
| β-CATENIN | Rbt | Cell sig 8480 | 1:500 |
| SLUG | Rbt | Cell sig 9585 | 1:500 |
| OCT4 | Ms | Sc5279 | 1:500 |
| SOX2 | Rbt | Ab97959 | 1:500 |
| NANOG | Ms | Ab62734 | 1:500 |

**Supplementary Figure S1.**
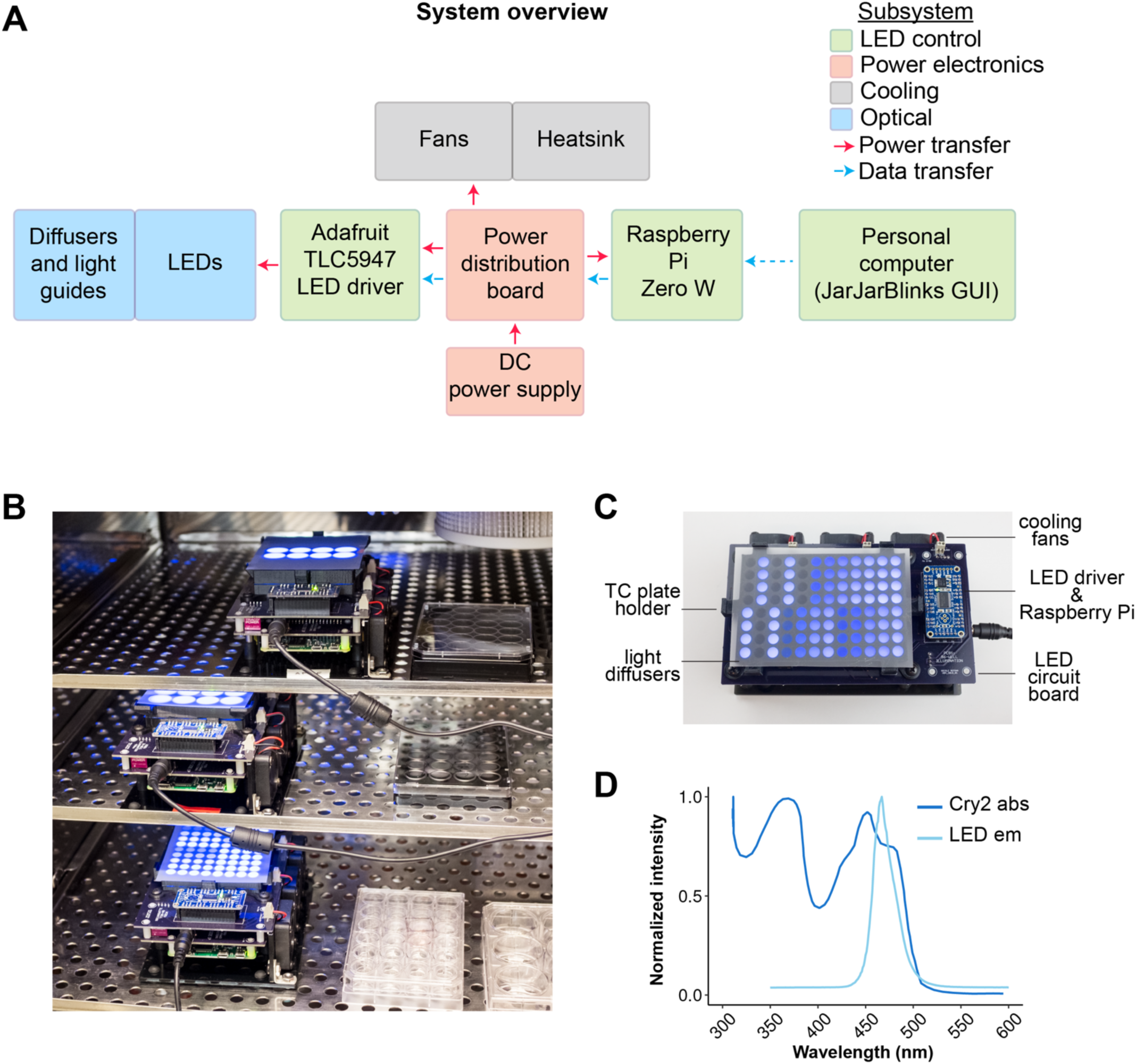
Overview of 24-well and 96-well LAVA devices. **a)** System block diagram of LAVA device. **b)** Image of 24-well and 96-well LAVA devices in TC incubator. **c)** Image of assembled 96-well LAVA board, with optical, cooling, and electronic subsystems highlighted. **d)** Emission spectrum of 470 nm blue LEDs match the absorption spectrum of Cry2. Cry2 spectrum adapted from reference (Banerjee et al. 2007).

**Supplementary Figure S2.**
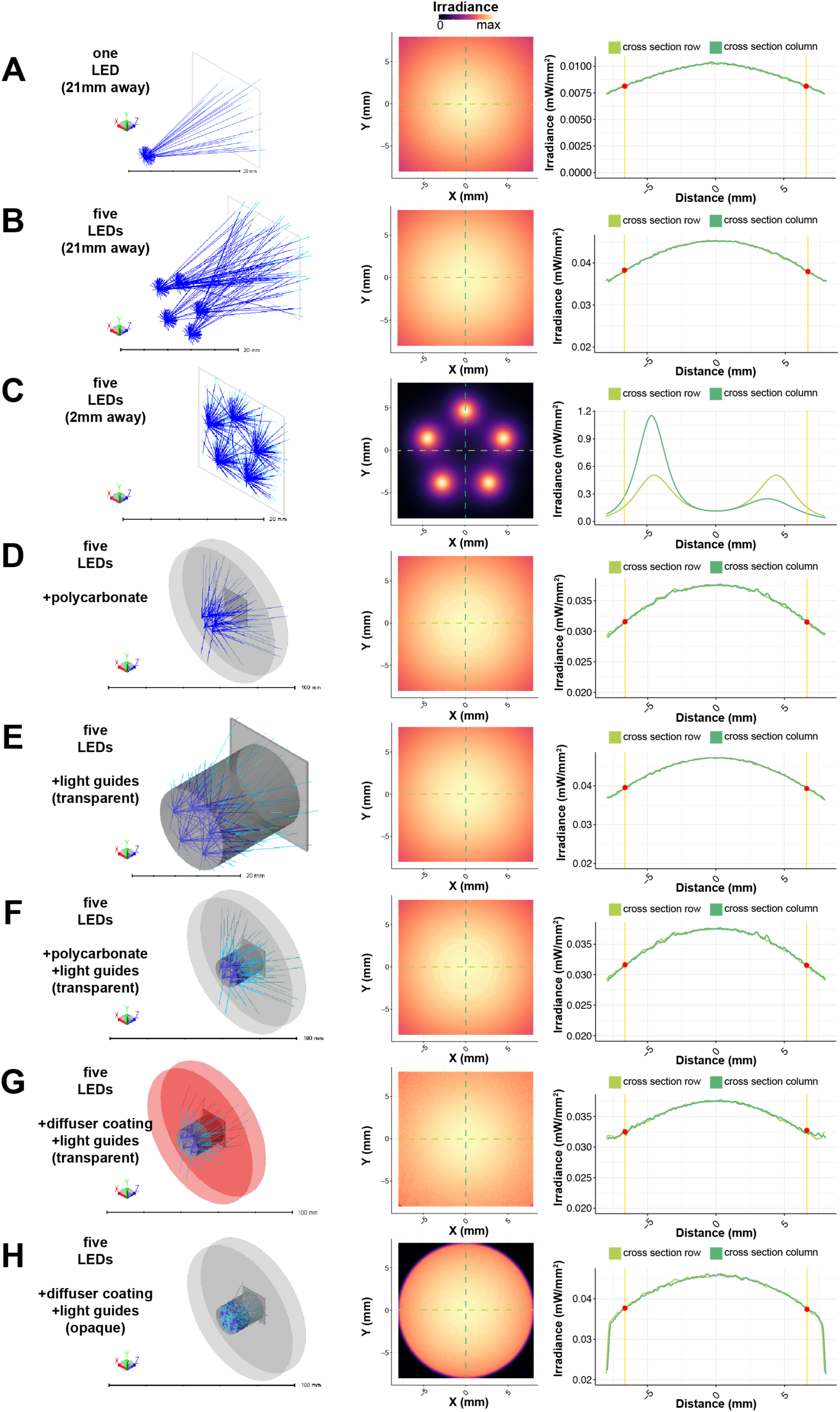
In silico validation of Zemax ray tracing model. Schematic of LED configuration (left), modeling result at detector plane (middle), and column and row cross-sections (right) with well edge of 24-well plate indicated with red points. **a)** Single LED illuminating detector 21mm away. **b)** Five LEDs, distributed along 1cm diameter circle, illuminating detector 21mm away. **c)** Five LEDs illuminating detector 2mm away. **d)** Five LEDs illuminating detector 21mm away through two 0.01” thick sheets of polycarbonate. **e)** Five LEDs illuminating detector 21mm away through two 10mm transparent light guides. **f)** Five LEDs illuminating detector 21mm away through two 0.01” thick sheets of polycarbonate and two 10mm transparent light guides. **g)** Five LEDs illuminating detector 21mm away through two 0.01” thick sheets of polycarbonate with 80° diffuser coating and two 10mm transparent light guides. **h)** Five LEDs illuminating detector 21mm away through two uncoated 0.01” thick sheets of polycarbonate and two 10mm reflective light guides with Lambertian scattering.

**Supplementary Figure S3.**
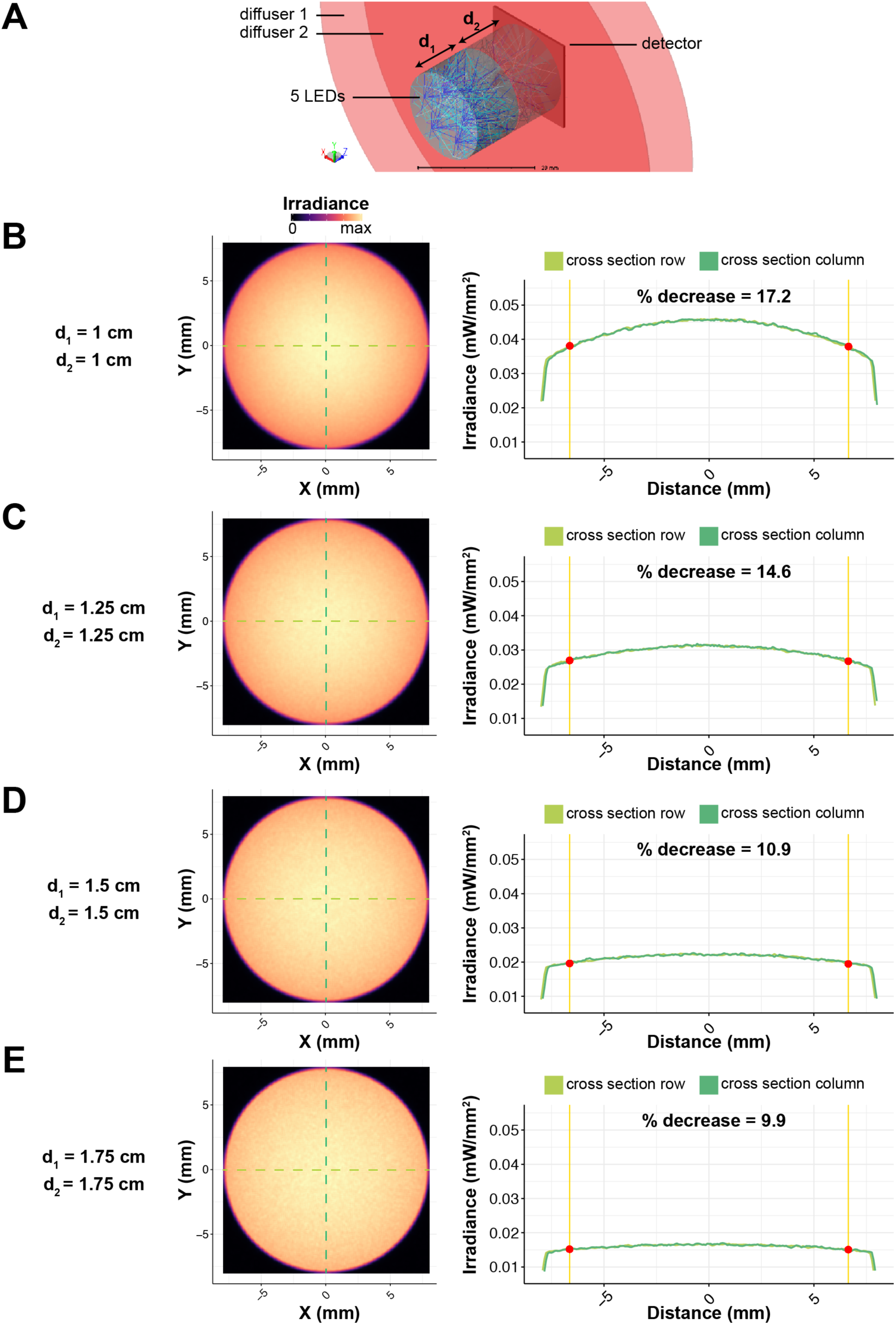
Results of Zemax modeling at variable light guide thicknesses, d_1_ and d_2_. **a)** Schematic of modeling setup. Five LEDs illuminate detector through two 0.01” thick sheets of polycarbonate with 80° diffuser coating (red) and two reflective light guides with Lambertian scattering (grey cylinder). **b-e)** Modeling results at indicated values of d_1_, d_2_. Image at detector plane (left) and column and row cross-sections (right) with well edge of 24-well plate indicated with red points show improved illumination uniformity at expense of light intensity with increasing light guide thickness.

**Supplementary Figure S4.**
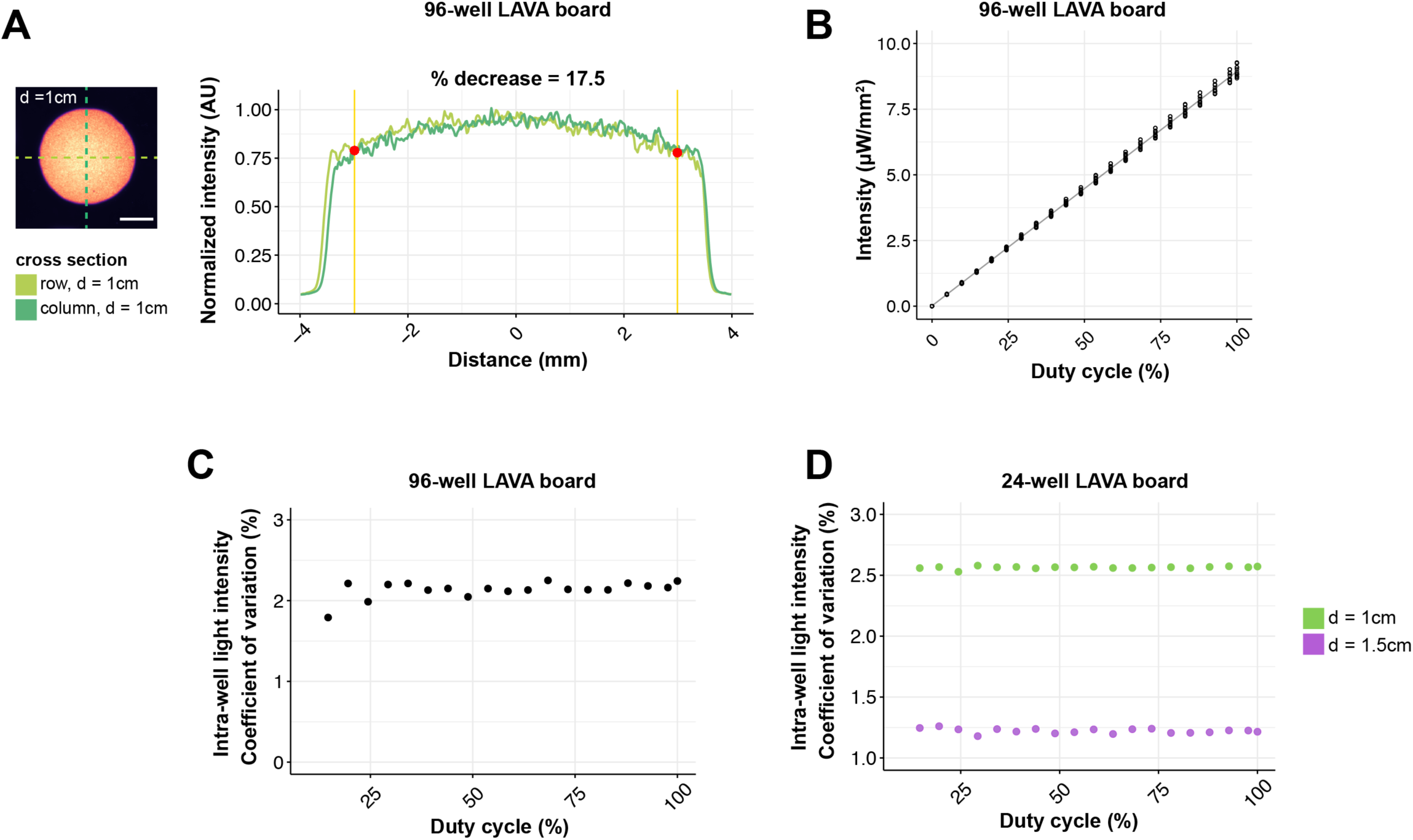
Intensity characterization of 24-well and 96-well LAVA boards. **a)** Brightfield images of well (left) and graph of intensity linescans along indicated cross-sections (right) characterize the intensity uniformity of the 96-well LAVA board when d = 1 cm. Percent decrease is calculated between intensity at center of well and intensity at highlighted red point, which indicates location of well edge of a 96-well culture plate. Graph shows mean normalized intensity over 2 independent wells. Scale bar 2.5 mm. **c)** Measured light intensity in response to the programmed duty cycle of the LED pulse-width modulation signal. Graph shows measured intensity from wells of a 96-well LAVA board and curve fit to a linear regression model. **d)** Coefficient of variation of light intensity between the 24 independent light channels of a 96-well LAVA device measured at different programmed intensities. **e)** Coefficient of variation of light intensity between the 24 independent light channels of a 24-well LAVA device measured at different programmed intensities. Green points correspond to optical configuration with d = 1 cm, violet points show optical configuration with d = 1.5 cm.

**Supplementary Figure S5.**
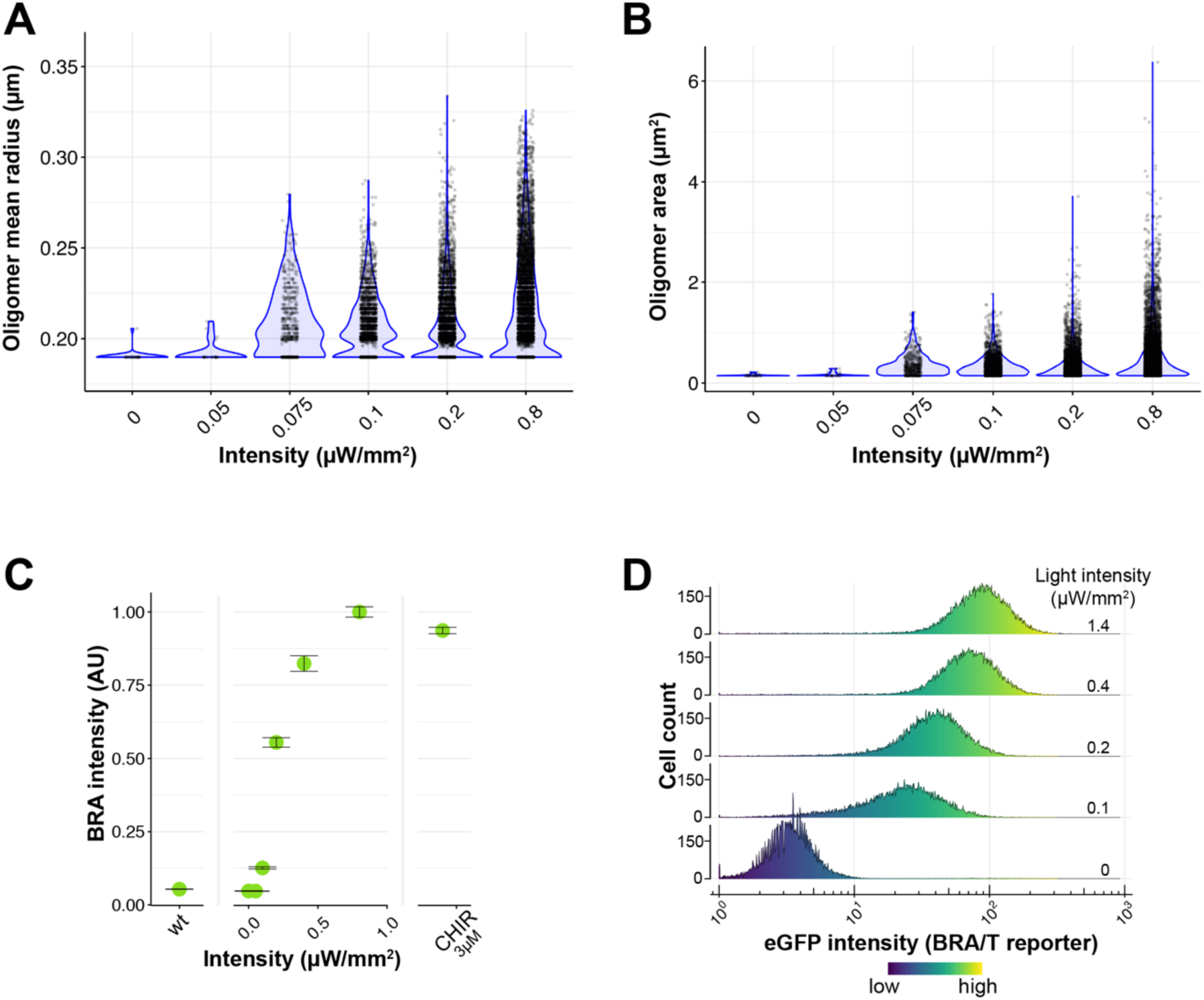
LRP6 oligomer size and BRA expression in optoWnt cells is dose-responsive to light intensity. **a-b)** Quantification of LRP6 oligomer size from immunostaining of optoWnt hESCs illuminated at indicated light intensities for 1 hr, Each point represents an LRP6 oligomer, with > 100 cells analyzed per condition. **c)** Quantification BRA immunostaining in response to increasing light intensity after 24 hr illumination or 3µM CHIR treatment. Average BRA intensity per hESC was calculated for each biological replicate. Graph shows mean of biological replicates ± 1 s.d., n = 3 replicates. **d)** Flow cytometry histograms of optoWnt hESCs expressing eGFP reporter for BRA/T after 24 hr illumination at varying light intensities. Graph shows sum of n = 3 replicates.

**Supplementary Figure S6.**
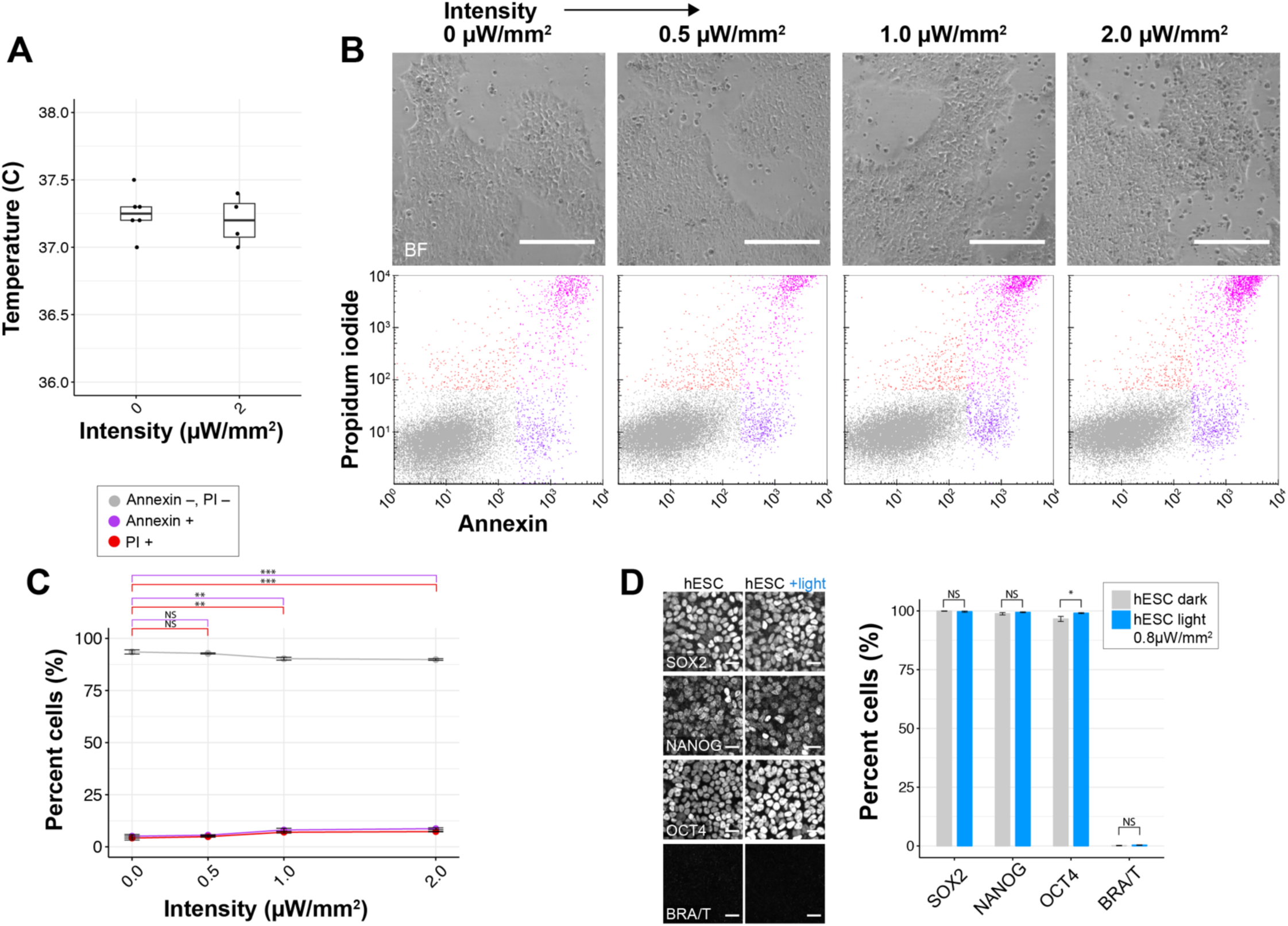
Phototoxicity during continuous optogenetic stimulation of hESC cultures. **a)** Temperature of media after 24 hrs of continuous illumination **b)** Brightfield images (top) of live wild-type hESC cultures illuminated at indicated light intensities for 48 hrs and flow cytometry results for Annexin V and propidium iodide (PI) stain. Scale bar 250 µm. **c)** Quantification of apoptosis marker Annexin V and dead cell stain PI shows a significant increase in apoptosis and cell death above 1µW/mm^2^ illumination intensity (p_A_=0.002, p_PI_=0.001 at 0 vs. 1 µW/mm^2^ and p_A_=0.0005, p_PI_ = 0.0005 at 0 vs. 2 µW/mm^2^). No difference was observed between 0 and 0.5µW/mm^2^ (p_A_=0.78, p_PI_=0.50). ANOVA followed by Tukey test. Graph shows mean ± 1 s.d., n = 3 biological replicates. **d)** Representative fluorescence images (left) and quantification (right) of wild-type hESCs stained for cell fate markers SOX2, NANOG, OCT4, and BRA after 48 hrs illumination at 0.8 µW/mm^2^. Student’s t-test (two-tail). Graph shows mean ± 1 s.d., n = 3 biological replicates. Scale bar 25 µm.

**Supplementary Figure S7.**
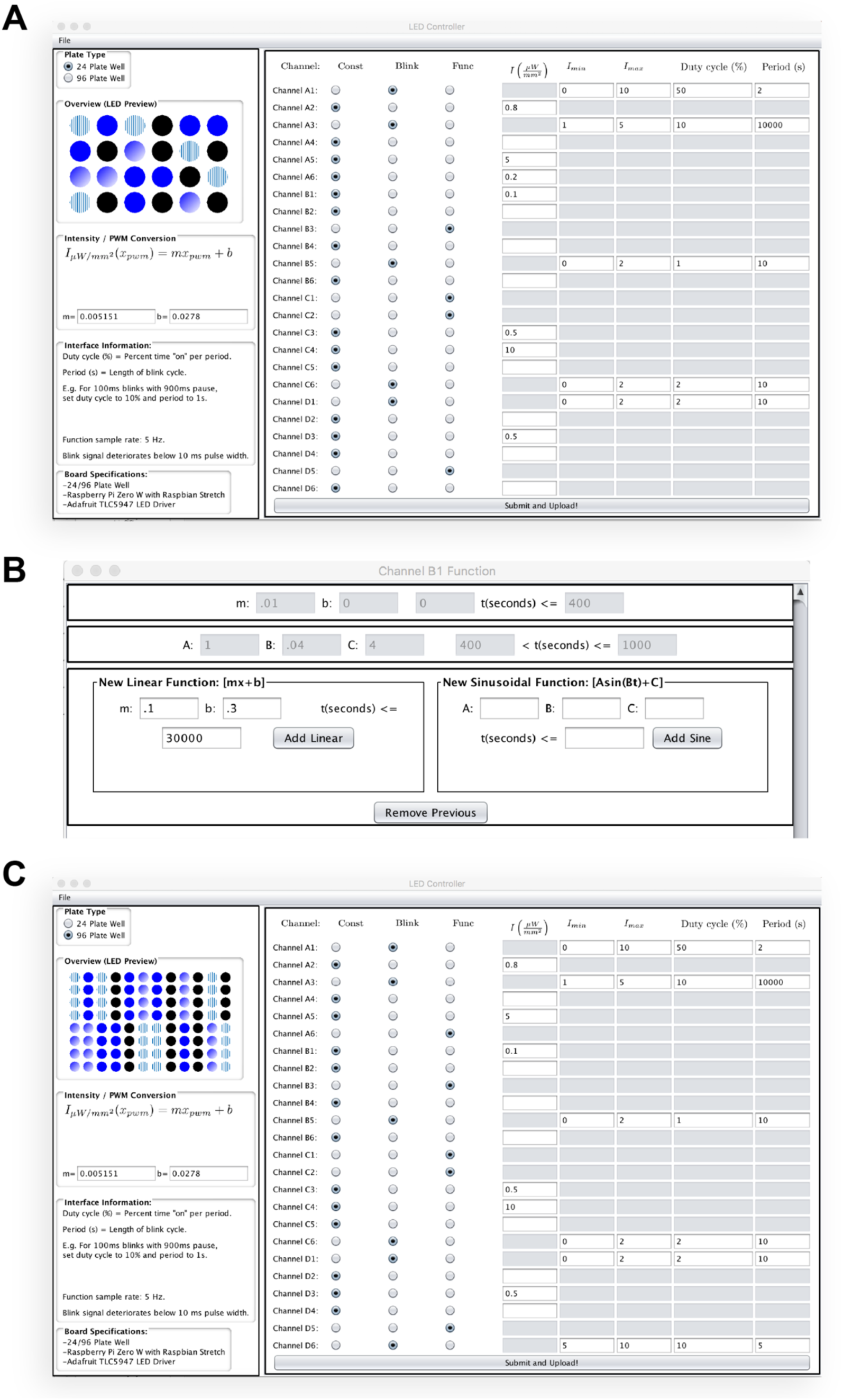
Screenshot of GUI for LAVA board control. **a)** Screenshot of GUI under 24-well TC plate setting **b)** Screenshot of diaolog box for input of time-varying light patterns as piecewise functions **c)** Screenshot of GUI under 96-well TC plate setting.

**Supplementary Figure S8.**
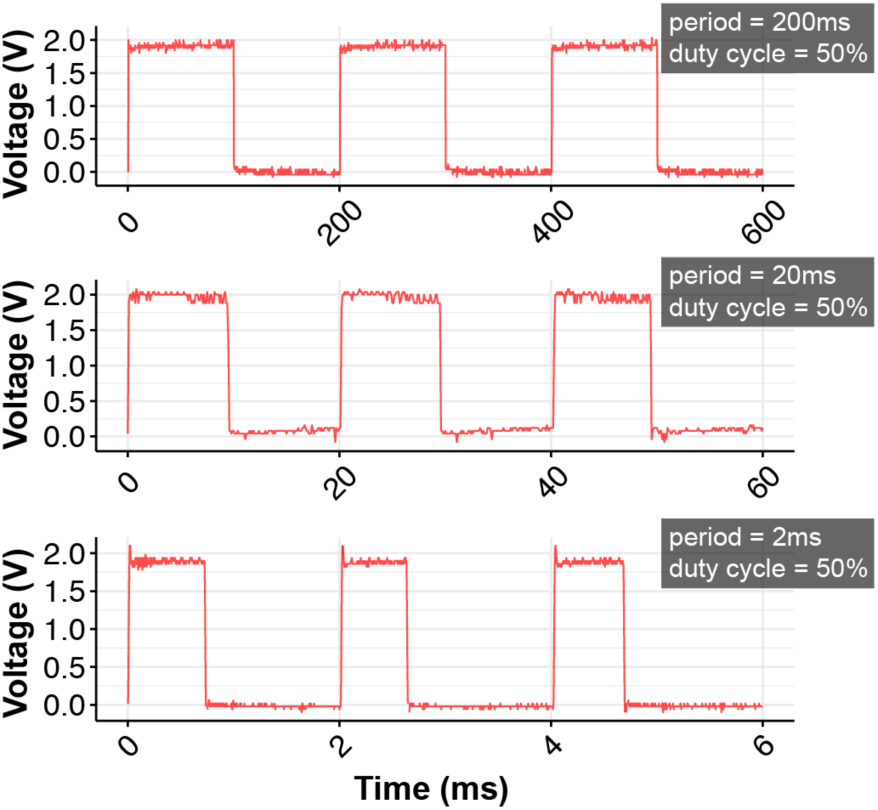
Measured illumination intensity during programmed blink sequences show signal inaccuracy at 1 ms pulses. Voltage signal from power meter measured with oscilloscope and is proportional to irradiance.

**Supplementary Figure S9.**
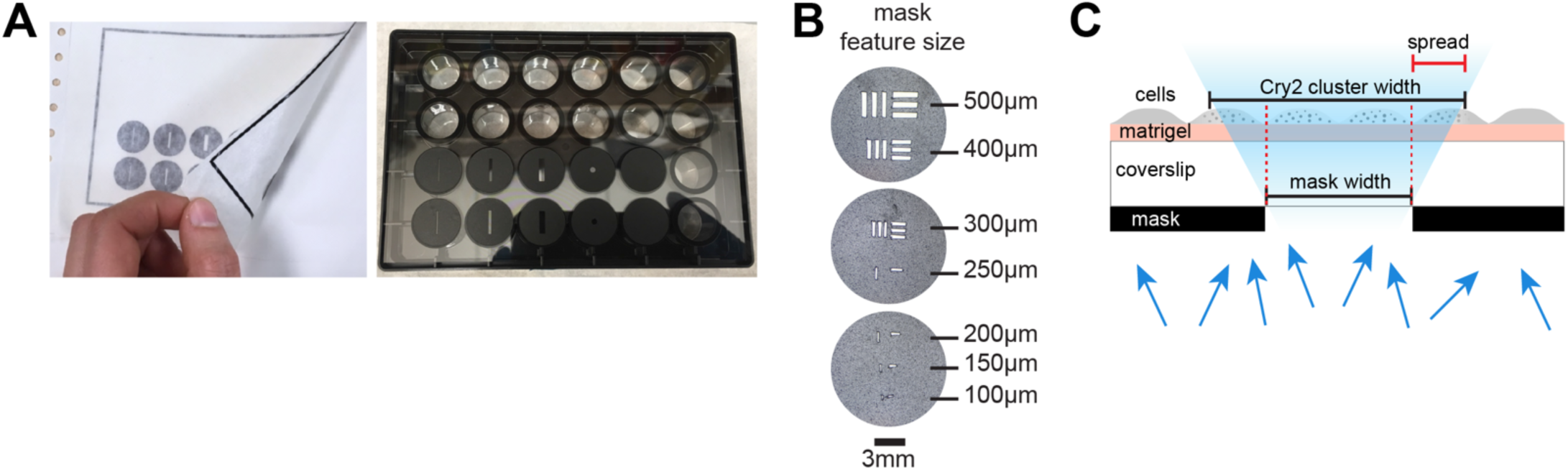
Dye-cut photomask enables spatial control of illumination **a)** Images of adhesive die-cut masks applied using transfer tape (top) onto 24-well TC plate (bottom). **b)** Brightfield images of die-cut mask illustrate resolution limit of cutter. Scale bar 3 mm. **c)** Schematic of light scattering from photomask.

### Description of additional supplementary files

File name: SupplementaryVideo1

Description:

**Supplementary Video 1: 24-well LAVA board displaying programmed light sequences**.

## REFERENCES

Acker, L. et al., 2016. FEF inactivation with improved optogenetic methods. Proceedings of the National Academy of Sciences of the United States of America, 113(46), pp.E7297–E7306.

Allen, B.D., Singer, A.C. & Boyden, E.S., 2015. Principles of designing interpretable optogenetic behavior experiments. Learning & memory (Cold Spring Harbor, N.Y.), 22(4), pp.232–238.

Arnold, S.J. & Robertson, E.J., 2009. Making a commitment: cell lineage allocation and axis patterning in the early mouse embryo. Nat. Rev. Mol. Cell Biol., 10(2), pp.91–103.

Arnold, S.J. et al., 2000. Brachyury is a target gene of the Wnt/β-catenin signaling pathway. Mechanisms of Development, 91(1-2), pp.249–258.

Aulehla, A. et al., 2003. Wnt3a plays a major role in the segmentation clock controlling somitogenesis. Dev. Cell, 4(3), pp.395–406.

Banerjee, R. et al., 2007. The signaling state of Arabidopsis cryptochrome 2 contains flavin semiquinone. J. Biol. Chem., 282(20), pp.14916–14922.

Bao, X. et al., 2016. Long-term self-renewing human epicardial cells generated from pluripotent stem cells under defined xeno-free conditions. Nature Biomedical Engineering, 1(1), p.0003.

Benedetti, L. et al., 2018. Light-activated protein interaction with high spatial subcellular confinement. Proceedings of the National Academy of Sciences of the United States of America, 115(10), pp.E2238–E2245.

Bernardo, A.S. et al., 2011. BRACHYURY and CDX2 mediate BMP-induced differentiation of human and mouse pluripotent stem cells into embryonic and extraembryonic lineages. Cell Stem Cell, 9(2), pp.144–155.

Blauwkamp, T.A. et al., 2012. Endogenous Wnt signalling in human embryonic stem cells generates an equilibrium of distinct lineage-specified progenitors. Nat Commun, 3(1), p.1070.

Boyden, E.S. et al., 2005. Millisecond-timescale, genetically targeted optical control of neural activity. Nature Neuroscience, 8(9), pp.1263–1268.

Bugaj, L.J. et al., 2018. Cancer mutations and targeted drugs can disrupt dynamic signal encoding by the Ras-Erk pathway. Science, 361(6405), p.eaao3048.

Bugaj, L.J. et al., 2013. Optogenetic protein clustering and signaling activation in mammalian cells. Nat. Methods, 10(3), pp.249–252.

Bugaj, L.J. et al., 2015. Regulation of endogenous transmembrane receptors through optogenetic Cry2 clustering. Nat Commun, 6, p.6898.

Carpenter, A.E. et al., 2006. CellProfiler: image analysis software for identifying and quantifying cell phenotypes. Genome biology, 7(10), p.R100.

Carrillo-Reid, L. et al., 2016. Imprinting and recalling cortical ensembles. Science, 353(6300), pp.691–694.

Čapek, D. et al., 2019. Light-activated Frizzled7 reveals a permissive role of non-canonical wnt signaling in mesendoderm cell migration. eLife., 8, p.1025.

Davidson, E.A., Basu, A.S. & Bayer, T.S., 2013. Programming microbes using pulse width modulation of optical signals. J. Mol. Biol., 425(22), pp.4161–4166.

Davidson, K.C. et al., 2012. Wnt/β-catenin signaling promotes differentiation, not self-renewal, of human embryonic stem cells and is repressed by Oct4. Proceedings of the National Academy of Sciences of the United States of America, 109(12), pp.4485–4490.

Duan, L. et al., 2017. Understanding CRY2 interactions for optical control of intracellular signaling. Nat Commun, 8(1), p.547.

Folch, A., 2016. Introduction to BioMEMS 1st ed., CRC Press.

Gerhardt, K.P. et al., 2016. An open-hardware platform for optogenetics and photobiology. Sci Rep, 6(1), p.35363.

Guglielmi, G. et al., 2015. An Optogenetic Method to Modulate Cell Contractility during Tissue Morphogenesis. Dev. Cell, 35(5), pp.646–660.

Guntas, G. et al., 2015. Engineering an improved light-induced dimer (iLID) for controlling the localization and activity of signaling proteins. Proceedings of the National Academy of Sciences of the United States of America, 112(1), pp.112–117.

Hannanta-Anan, P. & Chow, B.Y., 2016. Optogenetic Control of Calcium Oscillation Waveform Defines NFAT as an Integrator of Calcium Load. Cell Syst, 2(4), pp.283–288.

Hegemann, P. & Nagel, G., 2013. From channelrhodopsins to optogenetics. EMBO Molecular Medicine, 5(2), pp.173–176.

Hennemann, J. et al., 2018. Optogenetic Control by Pulsed Illumination. Chembiochem, 19(12), pp.1296–1304.

Hernandez, O. et al., 2016. Three-dimensional spatiotemporal focusing of holographic patterns. Nat Commun, 7(1), p.11928.

Huang, A. et al., 2017. Decoding temporal interpretation of the morphogen Bicoid in the early Drosophila embryo. eLife

Izquierdo, E., Quinkler, T. & De Renzis, S., 2018. Guided morphogenesis through optogenetic activation of Rho signalling during early Drosophila embryogenesis. Nat Commun, 9(1), p.2366.

Johnson, H.E. & Toettcher, J.E., 2018. Illuminating developmental biology with cellular optogenetics. Current opinion in biotechnology, 52, pp.42–48.

Johnson, H.E. & Toettcher, J.E., 2019. Signaling Dynamics Control Cell Fate in the Early Drosophila Embryo. Dev. Cell, 48(3), pp.361–370.e3.

Johnson, H.E. et al., 2017. The Spatiotemporal Limits of Developmental Erk Signaling. Dev. Cell, 40(2), pp.185–192.

Kainrath, S. et al., 2017. Green-Light-Induced Inactivation of Receptor Signaling Using Cobalamin-Binding Domains. Angew Chem, 56(16), pp.4608–4611.

Kempf, H. et al., 2016. Bulk cell density and Wnt/TGFbeta signalling regulate mesendodermal patterning of human pluripotent stem cells. Nat Commun, 7(1), p.13602.

Kiecker, C. & Niehrs, C., 2001. A morphogen gradient of Wnt/beta-catenin signalling regulates anteroposterior neural patterning in Xenopus. Development, 128(21), pp.4189–4201.

Kim, S.-E. et al., 2013. Wnt stabilization of β-catenin reveals principles for morphogen receptor-scaffold assemblies. Science, 340(6134), pp.867–870.

Kimura-Yoshida, C. et al., 2005. Canonical Wnt Signaling and Its Antagonist Regulate Anterior-Posterior Axis Polarization by Guiding Cell Migration in Mouse Visceral Endoderm. Developmental Cell, 9(5), pp.639–650.

Kirkeby, A. et al., 2012. Generation of regionally specified neural progenitors and functional neurons from human embryonic stem cells under defined conditions. Cell Rep, 1(6), pp.703–714.

Lee, J.M. et al., 2013. Switchable gene expression in Escherichia coli using a miniaturized photobioreactor. S.-H. Yun, ed. PLoS ONE, 8(1), p.e52382.

Levskaya, A. et al., 2009. Spatiotemporal control of cell signalling using a light-switchable protein interaction. Nature, 461(7266), pp.997–1001.

Li, V.S.W. et al., 2012. Wnt signaling through inhibition of β-catenin degradation in an intact Axin1 complex. Cell, 149(6), pp.1245–1256.

Lian, X. et al., 2012. Robust cardiomyocyte differentiation from human pluripotent stem cells via temporal modulation of canonical Wnt signaling. Proceedings of the National Academy of Sciences of the United States of America, 109(27), pp.E1848–57.

Lindsley, R.C. et al., 2006. Canonical Wnt signaling is required for development of embryonic stem cell-derived mesoderm. Development, 133(19), pp.3787–3796.

Liu, P. et al., 1999. Requirement for Wnt3 in vertebrate axis formation. Nat. Genet., 22(4), pp.361–365.

Luis, T.C. et al., 2011. Canonical wnt signaling regulates hematopoiesis in a dosage-dependent fashion. Cell Stem Cell, 9(4), pp.345–356.

Massey, J. et al., 2019. Synergy with TGFβ ligands switches WNT pathway dynamics from transient to sustained during human pluripotent cell differentiation. Proceedings of the National Academy of Sciences of the United States of America, 116(11), pp.4989–4998.

Müller, K., Zurbriggen, M.D. & Weber, W., 2014. Control of gene expression using a red- and far-red light-responsive bi-stable toggle switch. Nat Protoc, 9(3), pp.622–632.

Nagel, G. et al., 2003. Channelrhodopsin-2, a directly light-gated cation-selective membrane channel. Proceedings of the National Academy of Sciences of the United States of America, 100(24), pp.13940–13945.

Nikolenko, V., Poskanzer, K.E. & Yuste, R., 2007. Two-photon photostimulation and imaging of neural circuits. Nat. Methods, 4(11), pp.943–950.

Oates, A.C. et al., 2009. Quantitative approaches in developmental biology. Nat. Rev. Genet., 10(8), pp.517–530.

Olson, E.J. et al., 2014. Characterizing bacterial gene circuit dynamics with optically programmed gene expression signals. Nat. Methods, 11(4), pp.449–455.

Owen, S.F., Liu, M.H. & Kreitzer, A.C., 2019. Thermal constraints on in vivo optogenetic manipulations. Nat. Neurosci., pp.1–9.

Packer, A.M. et al., 2015. Simultaneous all-optical manipulation and recording of neural circuit activity with cellular resolution in vivo. Nat. Methods, 12(2), pp.140–146.

Packer, A.M., Roska, B. & Häusser, M., 2013. Targeting neurons and photons for optogenetics. Nat. Neurosci., 16(7), pp.805–815.

Papagiakoumou, E. et al., 2008. Patterned two-photon illumination by spatiotemporal shaping of ultrashort pulses. Optics Express, 16(26).

Papagiakoumou, E. et al., 2010. Scanless two-photon excitation of channelrhodopsin-2. Nat. Methods, 7(10), pp.848–854.

Pégard, N.C. et al., 2017. Three-dimensional scanless holographic optogenetics with temporal focusing (3D-SHOT). Nat Commun, 8(1), p.1228.

Prakash, R. et al., 2012. Two-photon optogenetic toolbox for fast inhibition, excitation and bistable modulation. Nat. Methods, 9(12), pp.1171–1179.

Repina, N.A. et al., 2017. At Light Speed: Advances in Optogenetic Systems for Regulating Cell Signaling and Behavior. Annual review of chemical and biomolecular engineering, 8, pp.13– 39.

Repina, N.A. et al., 2019. Optogenetic control of Wnt signaling for modeling early embryogenic patterning with human pluripotent stem cells. bioRxiv, 8, p.665695.

Richter, F. et al., 2015. Upgrading a microplate reader for photobiology and all-optical experiments. Photochem. Photobiol. Sci. 14(2), pp.270–279.

Rivera-Pérez, J.A. & Magnuson, T., 2005. Primitive streak formation in mice is preceded by localized activation of Brachyury and Wnt3. Developmental Biology, 288(2), pp.363–371.

Sako, K. et al., 2016. Optogenetic Control of Nodal Signaling Reveals a Temporal Pattern of Nodal Signaling Regulating Cell Fate Specification during Gastrulation. Cell Rep, 16(3), pp.866–877.

Schindelin, J. et al., 2012. Fiji: an open-source platform for biological-image analysis. Nat. Methods, 9(7), pp.676–682.

Shao, J. et al., 2018. Synthetic far-red light-mediated CRISPR-dCas9 device for inducing functional neuronal differentiation. Proceedings of the National Academy of Sciences of the United States of America, 8, p.201802448.

Sonnen, K.F. et al., 2018. Modulation of Phase Shift between Wnt and Notch Signaling Oscillations Controls Mesoderm Segmentation. Cell, 172(5), pp.1079–1090.e12.

Strickland, D. et al., 2012. TULIPs: tunable, light-controlled interacting protein tags for cell biology. Nat. Methods, 9(4), pp.379–384.

Sumi, T. et al., 2008. Defining early lineage specification of human embryonic stem cells by the orchestrated balance of canonical Wnt/beta-catenin, Activin/Nodal and BMP signaling. Development, 135(17), pp.2969–2979.

Thiery, J.P. & Sleeman, J.P., 2006. Complex networks orchestrate epithelial–mesenchymal transitions. Nat Rev Mol Cell Biol., 7(2), pp.131–142.

Thiery, J.P. et al., 2009. Epithelial-mesenchymal transitions in development and disease. Cell, 139(5), pp.871–890.

Thomson, J.A. et al., 1998. Embryonic stem cell lines derived from human blastocysts. Science., 282(5391), pp.1145–1147.

Toettcher, J.E. et al., 2011. Light-based feedback for controlling intracellular signaling dynamics. Nat. Methods, 8(10), pp.837–839.

Toettcher, J.E., Weiner, O.D. & Lim, W.A., 2013. Using optogenetics to interrogate the dynamic control of signal transmission by the Ras/Erk module. Cell, 155(6), pp.1422–1434.

Tucker, C.L., Vrana, J.D. & Kennedy, M.J., 2014. Tools for controlling protein interactions using light. Curr Protoc Cell Biol, 64.

Tyssowski, K.M. & Gray, J.M., 2019. Blue light induces neuronal-activity-regulated gene expression in the absence of optogenetic proteins. bioRxiv, 12(1), p.572370.

Wang, X. et al., 2010. Light-mediated activation reveals a key role for Rac in collective guidance of cell movement in vivo. Nat. Cell Biol., 12(6), pp.591–597.

Warmflash, A. et al., 2014. A method to recapitulate early embryonic spatial patterning in human embryonic stem cells. Nat. Methods, 11(8), pp.847–854.

Williams, M. et al., 2012. Mouse primitive streak forms in situ by initiation of epithelial to mesenchymal transition without migration of a cell population. Developmental Dynamics, 241(2), pp.270–283.

Yamaguchi, T.P. et al., 1999. T (Brachyury) is a direct target of Wnt3a during paraxial mesoderm specification. Genes Dev., 13(24), pp.3185–3190.

Yazawa, M. et al., 2009. Induction of protein-protein interactions in live cells using light. Nat Biotechnol., 27(10), pp.941–945.

Yizhar, O. et al., 2011. Optogenetics in neural systems. Neuron, 71(1), pp.9–34.

Yu, W. et al., 2008. Wnt signaling determines ventral spinal cord cell fates in a time-dependent manner. Development, 135(22), pp.3687–3696.

Zeng, L. et al., 1997. The mouse Fused locus encodes Axin, an inhibitor of the Wnt signaling pathway that regulates embryonic axis formation. Cell, 90(1), pp.181–192.

